# Mechanosensitivity of amoeboid cells crawling in 3D

**DOI:** 10.1101/2021.05.11.443058

**Authors:** Florian Gaertner, Patricia Reis-Rodrigues, Ingrid de Vries, Miroslav Hons, Juan Aguilera, Michael Riedl, Alexander Leithner, Jack Merrin, Vanessa Zheden, Walter Anton Kaufmann, Robert Hauschild, Michael Sixt

**Affiliations:** Institute of Science and Technology Austria, 3400 Klosterneuburg, Austria; BIOCEV, First Faculty of Medicine, Charles University, Vestec, Czech Republic

**Keywords:** Cell migration, Confinement, Mechanical load, Amoeboid motility, Actin cytoskeleton, Wiskott-Aldrich Syndrome protein, Leukocytes

## Abstract

Efficient immune-responses require migrating leukocytes to be in the right place at the right time. When crawling through the body amoeboid leukocytes must traverse complex three-dimensional tissue-landscapes obstructed by extracellular matrix and other cells, raising the question how motile cells adapt to mechanical loads to overcome these obstacles. Here we reveal the spatio-temporal configuration of cortical actin-networks rendering amoeboid cells mechanosensitive in three-dimensions, independent of adhesive interactions with the microenvironment. In response to compression, Wiskott-Aldrich syndrom protein (WASp) assembles into dot-like structures acting as nucleation sites for actin spikes that in turn push against the external load. High precision targeting of WASp to objects as delicate as collagen fibers allows the cell to locally and instantaneously deform its viscoelastic surrounding in order to generate space for forward locomotion. Such pushing forces are essential for fast and directed leukocyte migration in fibrous and cell-packed tissues such as skin and lymph nodes.

**In Brief:** WASp-driven actin spikes counter compressive loads of crowded tissue-landscapes.

## INTRODUCTION

Cells imbedded in tissues are tightly confined by complex three-dimensional (3D) microenvironments. These can be dense fibrillar networks as in mesenchymal interstitia like the dermis or cell-packed environments such as organ parenchyma or lymphatic tissues (Weigelin et al., 2016). Due to the limited availability of open space in such environments, cells frequently experience compressive loads from their surroundings and this is particularly relevant for cells that actively migrate.

Single-cell migration is an almost ubiquitous phenomenon in eukaryote biology and follows a continuum of biophysical strategies, with the mesenchymal mode on one pole and the amoeboid mode on the other (Friedl and Wolf, 2010). Amoeboid cells, such as metastatic cancer cells or leukocytes migrate much faster and are more autonomous from their extracellular environment, than mesenchymal (or epithelial) cells, and rather than remodeling their environment they adapt to it (Paluch et al., 2016). Survival and locomotion of amoeboid cells do not depend on adhesive ligands and they migrate efficiently even in ectopic environments or artificial materials (Lammermann et al., 2008; Reversat et al., 2020). The quick shape changes that amoeboid cells owe their name to (Amoeba, from Greek *amoibe,* meaning “change”) are entirely autonomous from the environment and the cellular envelope behaves as an active surface that rapidly deforms even when the cell floats in suspension (Barry and Bretscher, 2010).

These active adaptations of cell shape seem necessary to negotiate tissues because ameboid cells do not digest their environment in order to create a path. Instead, in obstructive environments with heterogeneous geometry they effectively find the path of least resistance and chose larger pores over small ones (Renkawitz et al., 2019). Whenever there is no other choice and the cell faces a small pore, it actively deforms its cell body and/or transiently dilates the pore in order to pass through (Pflicke and Sixt, 2009; Thiam et al., 2016). How ameboid cells generate forces underlying these processes is incompletely understood.

In animal cells most intracellular forces are generated by the actin cytoskeleton, which produces pulling forces via actomyosin contraction and pushing forces via actin polymerization. Both pulling and pushing are tightly regulated by mechanical feedback. In order to test the geometrical and mechanical features of its surrounding, mesenchymal cells probe their substrate by pulling on it (Plotnikov and Waterman, 2013). To do so they use their integrin-mediated adhesion sites as mechanosensitive organelles, which emit signals in response to traction force. Substrate-probing by pulling forces seems less relevant for leukocyte migration. Whenever they move in an adhesion-free mode there is clearly no opportunity to pull. And whenever leukocytes do use adhesion receptors to generate friction with the substrate, they minimize traction: They tune their integrins so that they just transmit the extremely small traction forces required to move the cell body forward (Hons et al., 2018).

Pushing forces are generated whenever actin filaments polymerize against the plasma membrane (Mogilner, 2006). The prototypic pushing structure is the lamellipodium, the flat actin protrusion that forms the leading front of most amoeboid and mesenchymal cells (Krause and Gautreau, 2014). Lamellipodial actin can adapt its protrusive force to the counter-force it experiences, making the tip of a cell mechanosensitive (Heinemann et al., 2011). However, lamellipodia are strictly two dimensional (2D) leaflets and the molecular machinery that drives actin polymerization at the lamellipodial tip does not support growth in the third dimension (Fritz-Laylin et al., 2018; Fritz-Laylin et al., 2017). Hence, lamellipodia can protrude into open spaces, but they are insufficient to create the active surface of an amoeboid cell that adapts and deforms in 3D. In order to do that, the cell requires both, mechanical sensitivity and activity across its whole cortex and not only at its lamellipodial tips. Here we use dendritic cells (DCs) and T cells as model systems to address how cells respond to geometrical constraints of their environment. We investigate how these constraints are met by the molecular machinery that drives actin polymerization across the cellular cortex.

## RESULTS

### Actin spikes form in response to mechanical loads of restrictive environments

To study how amoeboid cells respond to compressive loads, we observed the migratory behavior of dendritic cells (DCs) under a patch of non-degradable agarose cast on a serum-coated coverslip (Hons et al., 2018) (**Figure 1A**). In this reductionist approach, no pre-formed space is available and the highly flattened cells need to actively lift and deform the agarose to create sufficient space between coverslip and agarose (**Figure 1A**) (Laevsky and Knecht, 2003). To precisely control the mechanical load that the cells have to counter, we adjusted the stiffnesses of the agarose around the range of cellular stiffness as measured by atomic force microscopy (AFM) (Blumenthal et al., 2020; Guimarães et al., 2020) (**Figures 1A, S1A**). With increasing load, the mean migratory speed declined (**Figures 1B,C, S1B**) and the cells shifted from continuous locomotion to a stop and go pattern where cells collapsed and stalled after short stretches of movement (**Figure 1D, Movie S1**). Notably, time spent in arrest rose only slightly from 2.5 kPa (below cellular stiffness) to 10 kPa (slightly exceeding cellular stiffness) but drastically at 17.5 kPa (**Figure 1D**). Morphometric analysis of average cell shape revealed that at low (2.5 kPa) and intermediate (10 kPa) load, DCs adopted an elongated shape with a broad leading edge and a slender cell body (**Figure 1E**, **Movie S1**). Only under high load (17.5 kPa), average cell shape became substantially broadened (**Figures 1E,F, S1C**). These data indicate that DCs can successfully counter restrictive environments of stiffnesses that are in the range of cellular stiffness but frequently collapse when external load becomes exceedingly high. The cortical actin cytoskeleton largely determines the cells’ mechanical properties (Chugh et al., 2017). To obtain a detailed map of actin distribution in DCs migrating under agarose we generated DCs expressing LifeAct-eGFP (see Methods). When migrating under agarose (10 kPa) and imaged with high-speed confocal microscopy, DCs displayed small actin-rich foci that were scattered across the cell area, with peak intensities in the cell body (**Figure 1G, Movie S2**). Notably, actin spikes were restricted to regions of cellular confinement (**Figure 1G, Movie S2).**DCs migrating under non-adhesive conditions display a substantial retrograde flow of actin as both cortical and lamellipodial actin slide backward in relation to the substrate (Renkawitz et al., 2009). Actin foci evolved when cells migrated between (inert) agarose and passivated coverslips, demonstrating their independence of adhesive interactions with the substrate (**Figure 1G, Movie S2**). Actin foci moved together with this bulk actin flow, indicating linkage of the foci to the rest of the actin cortex (**Figures 1G, S1D**). Over time, many actin foci evolved into elongated stripes (**Figures 1H**, **S1E, Movie S2**) and confocal z scans as well as correlated light and scanning electron microscopy (CLEM) indicated that these stripes corresponded to ridge- or spike-like surface structures that grow normal to the imaging plane and thus protrude into the agarose (**Figures 1H,I, S1F**). Ultrastructural analysis of serial sections suggested that actin-spikes generated pushing forces between the cell and the agarose overlay: when spikes where located on top of the nucleus, the protrusions into the agarose overlay were mirrored by an indentation in the nuclear lamina (**Figure 1J**).

**Figure 1.**
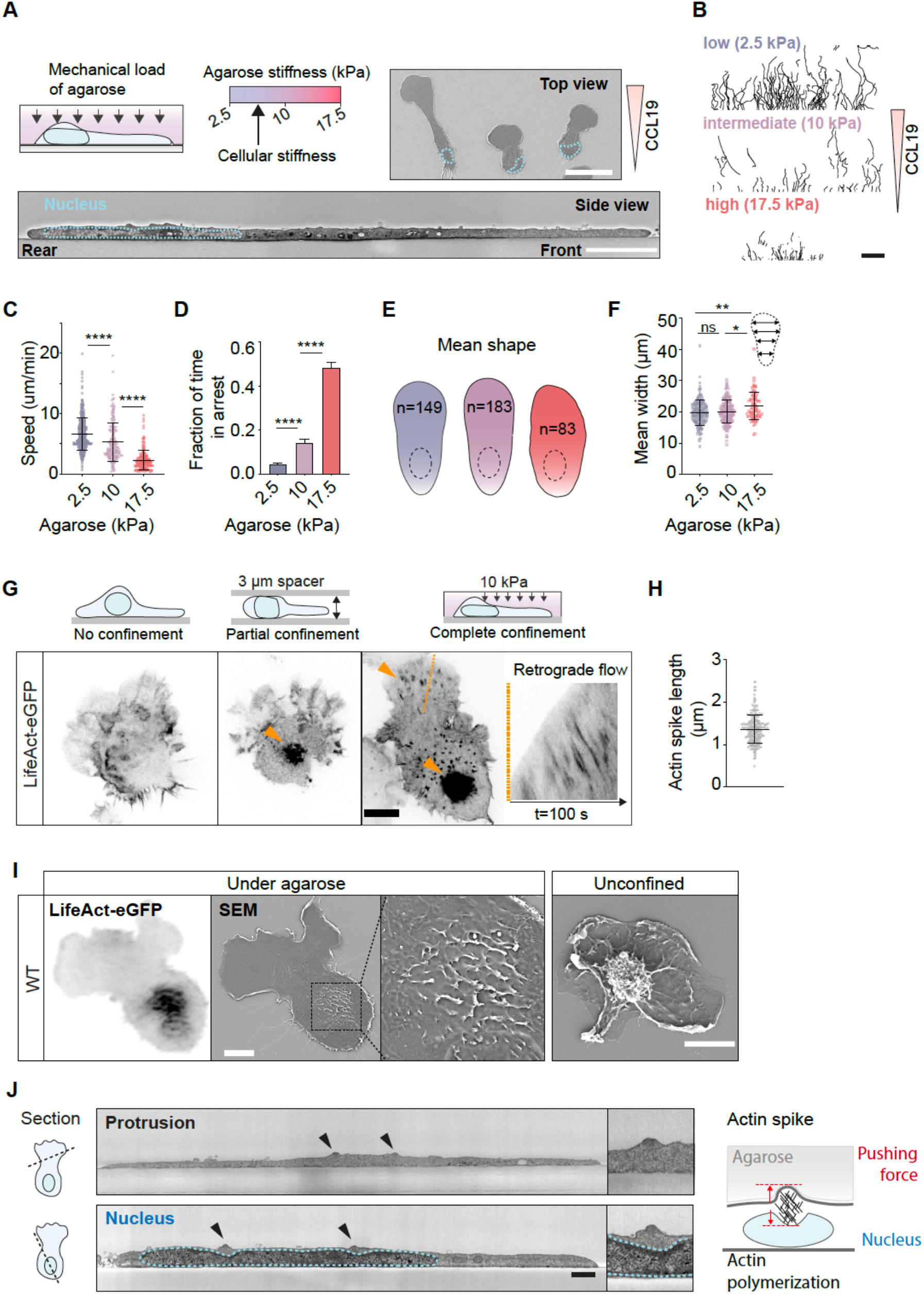
Actin spikes form in response to mechanical loads of restrictive environments. (**A**) Confinement of dendritic cells (DCs) under agarose. Agarose stiffness is adjusted around cellular stiffness (also see Figure S1A). Top view: SEM; scale bar=50 μm. Side view: SEM image of a sagittal section; scale bar=5 μm. (**B**) Plot of single cell tracks from representative experiments (△t=40 min); scale bar=200 μm. (**C**) Mean track speed. Each data point represents one track. 2.5 kPa: n=407 pooled from 6 experiments; 10 kPa: n=182 pooled from 4 experiments; 17.5 kPa: n=205 pooled from 4 experiments; Kruskal-Wallis/Dunn’s multiple comparisons test; ****: p<0.0001. Also see (Figure S1B). (**D**) Fraction of time in arrest was extracted from tracks in (C); Mean+SEM; Kruskal-Wallis/Dunn’s multiple comparisons test; ****: p<0.0001. (**E, F**) Mean shape of cell contours (E). Contours were further analyzed for mean cell width along the centerline (F). Kruskal-Wallis/Dunn’s multiple comparisons test; *: p=0.0189; **: p=0.0016. ns=not significant. (**G**) Actin spikes (LifeAct-eGFP) form in response to confinement. Representative images from spinning disc confocal microscopy movies of live cells on PMOXA-coated (non-adhesive) coverslips (confined cells). Unconfined cells adhere to PLL-coating. Kymograph shows retrograde actin flow. Notably, actin foci move with the bulk actin flow. (**H**) Actin spike length was quantified from confocal z-stacks (also see Figure S1F); n=246 pooled from 7 cells (Mean±SD). (**I**) Correlative light (epifluorescence) and (scanning) electron microscopy (CLEM). Note, LifeAct-eGFP signal correlates with spike and ridge-like protrusions. Right panel: SEM of unconfined cell. Scale bars=10 μm. (**J**) Ultrathin section analyzed by SEM. Upper panel: section across the leading edge. Lower panel: section across the cell body with nucleus. Notably, spikes (black arrow heads) form orthogonally to lamellipodia and indent the agarose. Mirrored indentations are visible in the nuclear lamina, indicating pushing forces generated by spikes (see magnified SEM images and graphical summary). Scale bar=1 μm. Also see Figure S1 and Movies S1 and S2.

Collectively these experiments suggest that cells confined under a restrictive overlay respond by forming actin spikes that exert vertical forces against the compressive load.

### WASp-driven actin spikes polymerize orthogonal to WAVE-driven lamellipodia

Formation and dynamics of the cortical actin cytoskeleton are mainly controlled by actin nucleators of the formin and Arp2/3 families (Bovellan et al., 2014). Thus, we tested how inhibition of formins and Arp2/3 with the small molecule inhibitors SMIFH2 and CK666 affect DC migration under agarose of intermediate (10kPa) stiffness. While formin inhibition mildly reduced migration speed (**Figure S2A**), Arp2/3 inhibition had a substantial effect with treated cells spending most of their time in arrest (**Figure S2B**). Notably, when mature DCs migrate in non-restrictive environments like cell-diameter sized microfluidic channels, inhibition of formins has similar effects on speed as it has under load while targeting Arp2/3 does not affect migration speed at all (Vargas et al., 2016). These findings suggest that Arp2/3 mediated actin nucleation becomes rate-limiting for DC locomotion whenever the cell has to counter external loads, while it is dispensable in non-restrictive environments. Hence, we focused on the Arp2/3 complex, which is activated by nucleation-promoting factors (NPFs) of the Wiskott-Aldrich Syndrome protein (WASp) and WASP-family verprolin-homologous protein (WAVE) families (Rotty et al., 2013). To test the contribution of these NPFs we analyzed DCs deficient in WASp or the WAVE complex subunit Hem1, which in hematopoietic cells is essential for the stability of the pentameric WAVE complex (Leithner et al., 2016a). Both mutants showed reduced migration speeds (**Figures 2A, S2C, Movie S3**) and increased time in arrest under agarose of intermediate stiffness (**Figure 2B**). In line with the fact that the WAVE complex exclusively drives horizontal forward protrusion at the very tip of the lamellipodium (Derivery and Gautreau, 2010; Graziano and Weiner, 2014), Hem1−/− DCs were devoid of lamellipodia and instead formed multiple, spiky protrusions at the leading edge (**Figure S2D**). Loss of lamellipodia renders DCs inefficient in exploring and navigating through complex environments (Leithner et al., 2016a), as indicated by prolonged times in arrest (**Figure 2B**). In contrast WASp−/− DCs formed normal lamellipodia (**Figure S2D**). Morphometric analysis showed that despite their spiky protrusions the mean overall cell body shape of Hem1−/− cells was comparable to WT cells, while WASp−/− cells were significantly broader (**Figure 2C**). Hence, regarding shape and migratory pattern, WASp−/− cells under intermediate-load conditions phenocopied WT cells under high-load conditions (**Figures 1B-F**).

**Figure 2.**
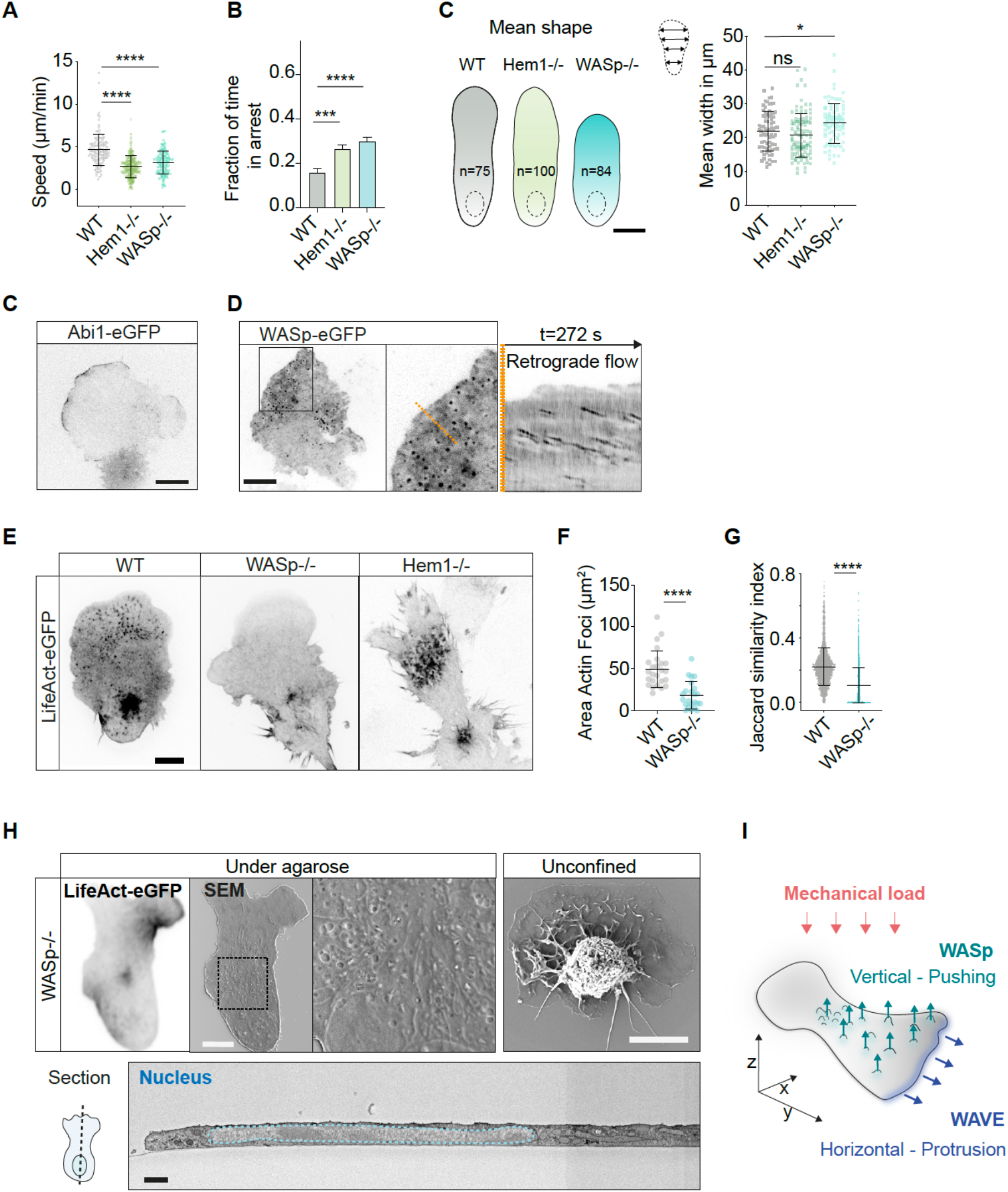
WASp-driven actin spikes polymerize orthogonal to WAVE-driven lamellipodia. (**A**) WASp−/− and Hem1−/− DCs were derived from bone marrow of transgenic mice. WT: n=118 pooled from 3 experiments; WASp−/−: n=175 pooled from 4 experiments; Hem1−/−: n=347 pooled from 6 experiments; Kruskal-Wallis/Dunn’s multiple comparisons test; ****: p<0.0001. Fraction of time in arrest was extracted from tracks; Mean+SEM; Mann Whitney test and Kruskal-Wallis/Dunn’s multiple comparisons test; ****: p<0.0001. (**B**) Mean shape of contours; WT: n=75; Hem1−/−: n=100; WASp−/−: n=84. Mean cell width along the centerline; Kruskal-Wallis/Dunn’s multiple comparisons test; (g) *: p=0.0491; ns=not significant. (**C,D**) TIRF microscopy of agarose-confined DCs expressing (C) Abi-eGFP (to label the WAVE complex) or (D) WASp-eGFP. Note, while Abi1-eGFP is restricted to the lamellipodial tip, WASp-eGFP forms dot-like structures scattered across the cell body and moving with the retrograde flow; resembling LifeAct-eGFP signal (see Figure 1G). Scale bar=10 μm. (**E**) WT, WASp−/− and Hem1−/− DCs under agarose (10kPa) on PMOXA-coated (non-adhesive) coverslips. Representative images from spinning disc confocal microscopy movies (LifeAct-eGFP) of live cells are shown (also see Movie S2). Notably, Hem1−/− DCs are devoid of lamellipodia, but form prominent actin foci at the leading edge and the cell body. Scale bar=10 μm. (**F, G**) Quantification of area (F) and dynamics (G) of actin foci in WT and WASp−/− cells. Jaccard index measures frame-to-frame overlap of segmented actin foci (1 = 100% similarity). (F) Area of actin foci. WT: n=25 cells; WASp−/−: n=23 cells. Mean±SD; Mann-Whitney; ****: p<0.0001. (G) WT: n=2144 frames pooled from 25 movies; WASp−/−: 2055 frames pooled from 23 movies. Mean±SD; Mann-Whitney; ****: p<0.0001. (**H**) Ultrastructural analysis of WASp−/− DCs using CLEM (upper panel; scale bars=10 μm) and SEM of an ultrathin section (lower panel; scale bar=1 μm). While unconfined WASp−/− cells morphologically resemble WT cells (Figure 1I), they do not form actin spike- and ridge-like structures upon confinement. (**I**) Graphical summary. Also see Figure S2 and Movie S3.

Next, we were interested in the spatio-temporal organization of WAVE and WASp-dependent actin networks under compressive loads. While the WAVE complex (detected with Abi1-eGFP) was strictly localized to the tip of lamellipodia (Leithner et al., 2016a), WASp (detected with WASp-eGFP) formed dots scattered across the cell surface, matching the distribution and flow-pattern of actin foci (**Figures 2C,D**). Deletion of WASp, but not the WAVE complex abrogated formation of actin foci and spikes as revealed by LifeAct-eGFP signal (**Figures 2E,F**), and the few actin foci remaining in WASp−/− DCs were transient and short-lived (**Figures 2G, S2E, Movie S3**). Ultrastructural analysis of serial sections and CLEM confirmed the morphological absence of spikes and ridges on the dorsal surface of WASp−/− DCs under confinement (**Figure 2H**), while there were no apparent morphological differences to WT cells in the absence of a restrictive overlay **(Figures 1I, 2H)**.

Together, these data reveal a characteristic organization of branched actin networks in 3D: while WAVE-driven actin nucleation powers the horizontal forward protrusive component of cellular locomotion, WASp-driven actin nucleation counters the mechanical load of the overlay (**Figure 2I**).

### Vertical pushing facilitates locomotion by deforming restrictive environments

We next used a cross-correlation approach to address if and how WASp-driven vertical pushing is embedded into the morphodynamic cycle of cellular locomotion. Acceleration of WT cells under agarose is initiated by leading-edge protrusion and elongation of the cell body (**Figure 3A, B, Movie S4**). Accordingly, cross-correlating speed with aspect ratio revealed a clear positive and instantaneous correlation (**Figures 3C, S3A**). Incorporating the number of cortical actin foci into the correlation showed that foci preceded acceleration by 1-2 min (**Figure 3C**). This sequence suggested that actin foci are a *prerequisite* to lift the load of the agarose, which then *facilitates* forward protrusion. We next measured nuclear-projected area as a proxy for how much a cell is able to lift the agarose: the nucleus of suspended DCs is approximately spherical, but naturally flattens with increasing confinement, making it a good indicator of cell height (**Figures 1A, S3B**). WT DCs showed steady fluctuations of nuclear-projected area when migrating under agarose (**Figure S3C**, **Movie S4**). Nuclear area was negatively cross-correlated with the appearance of actin foci on the surface area of the cell (**Figures 3C, D**), supporting the concept that vertical WASp-driven protrusions lift the agarose so that the nucleus can expand vertically. Accordingly, cross-correlations between (residual) actin foci, speed and projected nuclear area were blunted in WASp−/− DCs, while the speed to aspect ratio cross correlation was retained (**Figures 3F-H, Movie S4**). We next tested if the proposed morphodynamic cycle is indeed specific for migration in restrictive environments. We thus seeded DCs into PDMS-based (non-deformable) microfluidic devices with a constant height of 5 μm, which can be passaged without substrate deformation. Here, migration parameters of WASp−/− mutants were indistinguishable from WT cells (**Figure S3D, Movie S5**). To further challenge migration in non-deformable environments we reduced pillar spacings to 1.5 μm-sized constrictions **(Figure 3I)**. DCs squeezing through non-deformable constrictions recruited WASp and actin to sites of self-imposed compression (**Figure 3J, Movie S5**). We therefore tested if WASp-dependent pushing forces are required for self-deformation of the cell body and squeezing through non-deformable constrictions (Thiam et al., 2016). When encountering a constriction both WT and WASp−/− DCs arrested to squeeze their cell body through the pore. Importantly, mean time of passage of WASp−/− cells was not prolonged, but rather slightly reduced, indicating a negligible role of WASp-dependent pushing forces when countering non-deformable obstacles (**Figure 3K**). Together, we show that WASp-driven vertical pushing forces counter mechanical load and displace deformable microenvironments to generate space for the cell to enter into the protrusive phase of locomotion.

**Figure 3.**
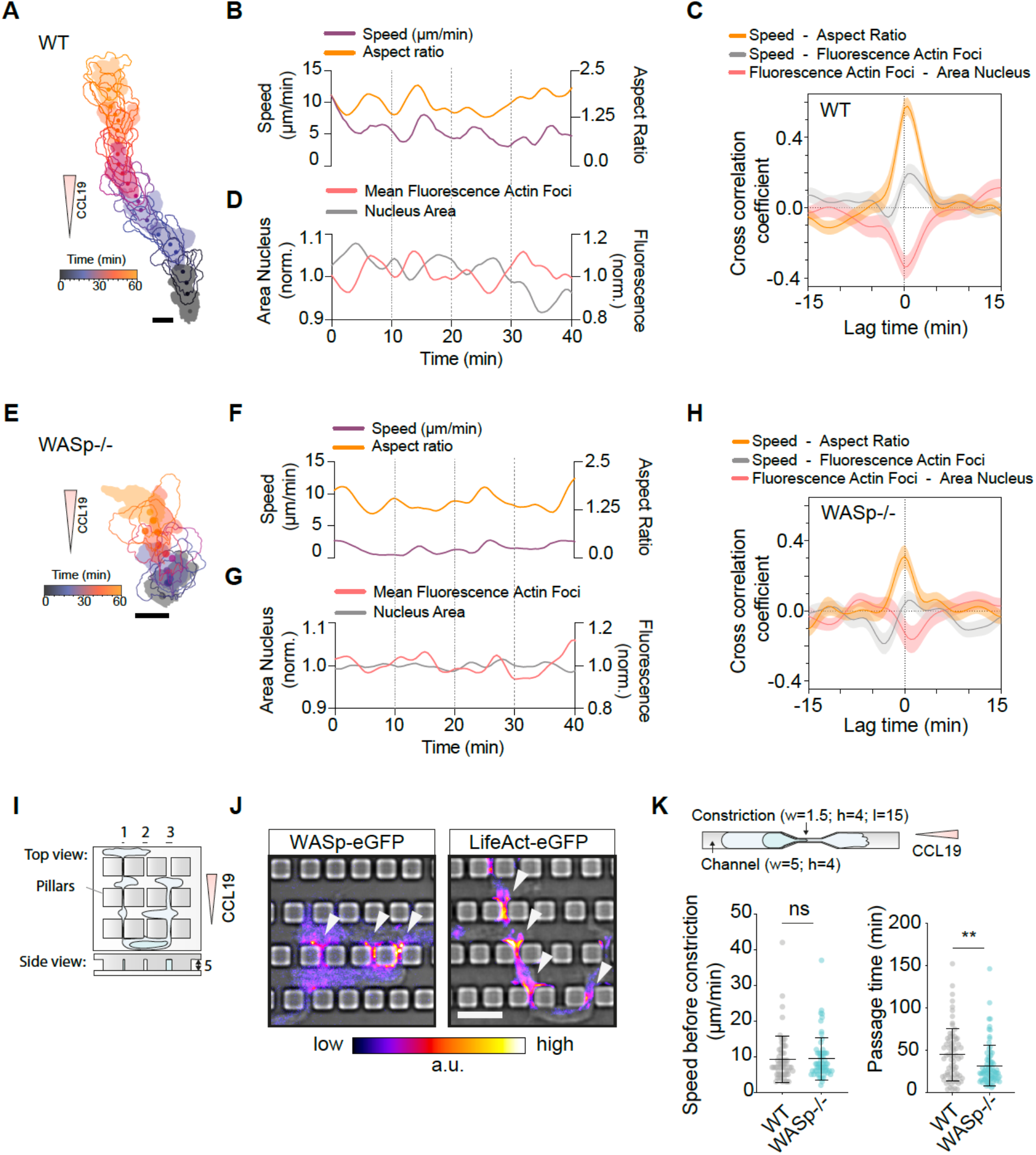
Vertical pushing facilitates locomotion by deforming restrictive environments. (**A**) Representative color-coded WT DC outlines (time). Scale bar=25 μm. (**B**) Smoothed time traces of speed and aspect ratio extracted from WT DC contours (epifluorescence movies (LifeAct-eGFP); frame rate=30s). (**C**) Cross-correlation between the (1) Cell Speed and Aspect Ratio; (2) Cell speed and Mean Fluorescence of Actin Foci and (3) Mean Fluorescence of Actin Foci and Area of Nucleus is shown (WT: n=25 cells; Mean±SEM). The positive lag time (gray curve) means an increase in Mean Fluorescence of Actin Foci precedes Cell Speed (also see Figure S3A). (**D**) Smoothed time traces of Area of Nucleus (normalized by the mean of all time points) and Mean Fluorescence of Actin Foci (normalized by the mean of all time points) of the same cell (see A, B). (**E**) Representative color-coded WASp−/− DC outlines (time). Scale bar=25 μm. (**F**) Smoothed time traces of speed and aspect ratio extracted from WASp−/− DC contours (epifluorescence movies (LifeAct-eGFP); frame rate=30s). (**G**) Smoothed time traces of Area of Nucleus (normalized by the mean of all time points) and Mean Fluorescence of Actin Foci (normalized by the mean of all time points) of the same cell (see E, F). (**H**) Cross-correlation analysis (see C) (WASp−/−: n=22-23 cells; Mean±SEM). (**I**) PDMS-based (non-deformable) microfluidic device with a constant height and constrictions. (**J**) Representative images of WASp-eGFP- and LifeAct-eGFP-expressing cells squeezing through constrictions (TIRF). Arrow heads indicate increased fluorescence at constrictions. Scale bar=10 μm. (**K**) Mean speed of cells migrating in straight channels and passage time of constriction. Schematic shows dimension of microfluidic device in μm. Also see Figure. S3 and Movies S4 and S5.

### WASp-driven actin polymerization is triggered by mechanical loading

We showed that WASp-driven vertical protrusions are (1) temporally correlated with the mechanical response and (2) required for countering external mechanical load. We next addressed causality and tested if actin foci form in response to a mechanical trigger or if they develop spontaneously.

We seeded LifeAct-eGFP expressing DCs on poly-l-lysine coated coverslips and locally pinched the cell body with the blunted tip of a microneedle (**Figure 4A**). Within seconds after indenting the cell, the actin reporter increased around the pipette tip of WT DCs, leaving behind a localized actin cloud (Gerard et al., 2014) (**Figure 4B, Movie S6**). This mechanically induced burst of actin polymerization was substantially decreased in WASp−/− DCs (**Figure 4C**). This finding supported a role of WASp in polymerizing actin in response to mechanical loading (**Figure 1G**). Leukocytes migrating within tissues actively bump into subcellular, usually submicron-sized obstacles such as collagen fibers that restrict their passage and therefore locally pinch the cell body. We engineered the geometric aspect of this scenario *in vitro* and confined cells onto inert substrates with a ridged topography matching the size of aligned collagen fibrils (**Figures 4D, S3A,B**). This setup allowed the precise localization of the fluorescent signals. We found that ridges of 600 nm height (and 250 nm width) induced actin foci that matched the pattern of ridges (**Figure 4E, Movie S6**). Notably, these foci exclusively formed on top of ridges and were virtually absent in the interjacent grooves (**Figures 4E, S4C, Movie S6**). This response was independent of adhesive interactions as actin foci formed efficiently on passivated substrates, where actin foci moved with the retrograde actin flow (**Figure 4E**). WASp-eGFP followed the same dynamic pattern, with sharply delineated dots precisely matching the ridge structure (**Figure 4F, Movie S6**). Importantly in WASp-deficient cells, the formation of actin foci on the ridges was virtually abrogated (**Figures 4E, S4C, Movie S6**). These data demonstrate that DCs show a highly localized response to mechanical indentation with rapid recruitment of WASp, which triggers polymerization of actin.

**Figure 4.**
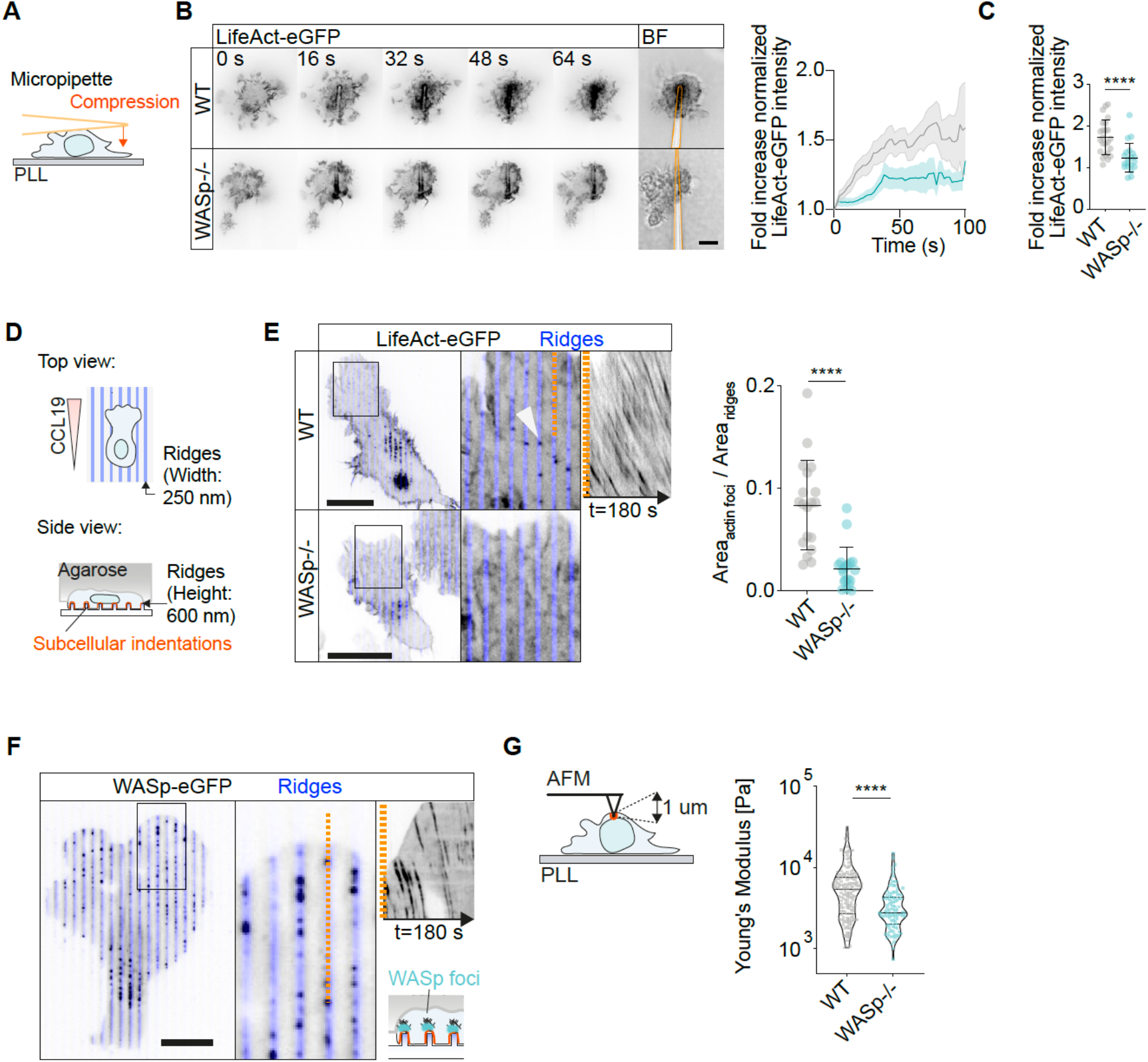
WASp-driven actin polymerization is triggered by mechanical loading. (**A**) DCs attached to poly-l-lysine (PLL)-coated coverslips were pinched from top with a microneedle (schematic). Micropipette was held in place during time-lapse recordings. Time traces (spinning disc confocal microscopy) of representative LifeAct-eGFP expressing DCs. (**B**) The last frame shows an overlay of brightfield images of the micropipette (highlighted in orange). Right panel: Time traces of LifeAct-eGFP signal at the site of micropipette indentation (normalized by mean LifeAct-eGFP signal of non-indented area). Intensity of first frame was set to 1. WT: n=24; WASp: n=20; Mean±SEM. Scale bar=10 μm. (**C**) Dot plot showing fold increase of Lifeact-eGFP intensity following indentation (last frame / first frame). Each data point is one indentation experiment (see B). Mean±SD; Mann-Whitney test; ****: p<0.0001. (**D**) Schematic shows DCs confined on inert nano-ridges with 2 μm pitch (PMOXA coating) (see Methods). (**E**) Representative WT and WASp−/− DC migrating on nano-ridges. Notably, actin foci formed with high precision on top of ridges but not in grooves (see Figure S4C). Arrow head indicates actin foci. Kymograph shows retrograde flow of actin foci. Fraction of ridges covered by actin foci (segmented in Ilastik) (also see Figure S4B). WT: n=19 cells; WASp−/−: n=19 cells; Mean±SD; Mann-Whitney test; ****: p<0.0001. Scale bar=20 μm. (**F**) Representative TIRF image from WASp-eGFP expressing DC migrating on nano-ridges (width: 1 μm; height: 250 nm; pitch: 2 μm). WASp-eGFP shows same dynamic pattern as actin foci (kymograph) with sharply delineated dots matching the ridge structure. Scale bar= 20 μm. Schematic of an AFM cantilever indenting a DC adherent to a PLL-coated coverslip (left). Using a Hertz contact mechanics model, the elastic modulus was estimated by fitting the force indentation curves up to 1 μm. Each data point represents one measurement. WT: n=160 from 37 cells (also see Figure S1A); WASp−/−: n=106 from 26 cells. For cellwise analysis see Figure S4D. Violin plot with Median±Quartiles; Mann-Whitney test; ****: p<0.0001. Also see Figure S4 and Movie S6.

Previous studies have shown that mechanical load can induce global stiffening of the cellular cortex, which is dependent on actin reorganization (Hu et al., 2019). To test if load-induced actin polymerization increases cortical stiffness in DCs we indented DCs immobilized on poly-l-lysine with the tip of an AFM cantilever and measured cellular stiffness. WASp-deficient DCs showed a significantly reduced cortical stiffness, indicating a crucial role of WASp in adapting cell mechanics to mechanical confinement (**Figures 4G, S4D**). In summary, we demonstrate that cells respond to mechanical load by locally polymerizing actin, a process requiring WASp.

### Deformation of collagen fibers is required for migration in fibrous environments

We next challenged our findings in physiological environments. To initiate adaptive immunity, DCs need to migrate from the site of antigen retrieval to the draining lymph node (LN) (Worbs et al., 2017), which requires fast transit through 3D tissues with complex geometry and molecular composition. DCs can reach the lymph node without proteolytically generating a path (Pflicke and Sixt, 2009), suggesting that they predominantly use a mechanical strategy. We therefore hypothesized that efficient migration in tissues requires WASp-driven actin to mechanically create space in dense 3D matrices. To test this, we first competitively injected WT and WASp−/− mature DCs into footpads of wildtype recipient mice to measure homing to draining popliteal LNs (**Figure S5A**). After 24h, recruitment of WASp−/− DCs was significantly reduced compared to WT controls, indicating a crucial role of WASp dependent actin polymerization *in vivo* and confirming previous reports by others (Bouma et al., 2007; de Noronha et al., 2005; Pulecio et al., 2008; Snapper et al., 2005).

To study the mechanism underlying impaired migration in fibrous 3D microenvironments in detail, we then used fibrillar collagen, the most abundant matrix scaffold of mammalian tissues, as a proxy for the interstitial matrix. Collagen gels were polymerized at densities resulting in pore sizes substantially smaller than the minimal cross-section of deformed DCs (Lang et al., 2015; Renkawitz et al., 2019; Thiam et al., 2016; Wolf et al., 2013). Consequently, chemotactic locomotion under these conditions should depend on the deformation of obstructive fibers (**Figure 5A**). Indeed, fast confocal imaging revealed that DCs locally displaced collagen fibers, thereby generating space for the passage of the cell body. Locally, the deformation of collagen fibers was accompanied by a burst in actin polymerization (**Figures 5B, C, Movie S7**). We then performed bulk measurements of large numbers of DCs and found that in low-density collagen gels, where pore sizes between fibers are sufficiently large to allow unconstrained locomotion, WASp−/− cells were not significantly slower than WT controls (**Figure 5D**). With increasing collagen gel densities (decreased pore size and increased stiffness), DCs were more reliant on displacing fibers to generate space (**Figure S5B**), became substantially slower (**Figure 5D**), and often arrested (**Figure 5E**), indicating that matrix deformation became a rate limiting parameter. Accordingly, under these conditions, WASp-deficient DCs were less efficient in recruiting actin to restrictive fibers **(Figures 5C, S5C**). Together, our data show that cells require Wasp-dependent pushing to mechanically create space in dense 3D matrices and provide a mechanistic explanation for impaired migration of WASp−/− DCs *in vivo* (**Figure S5A**).

**Figure 5.**
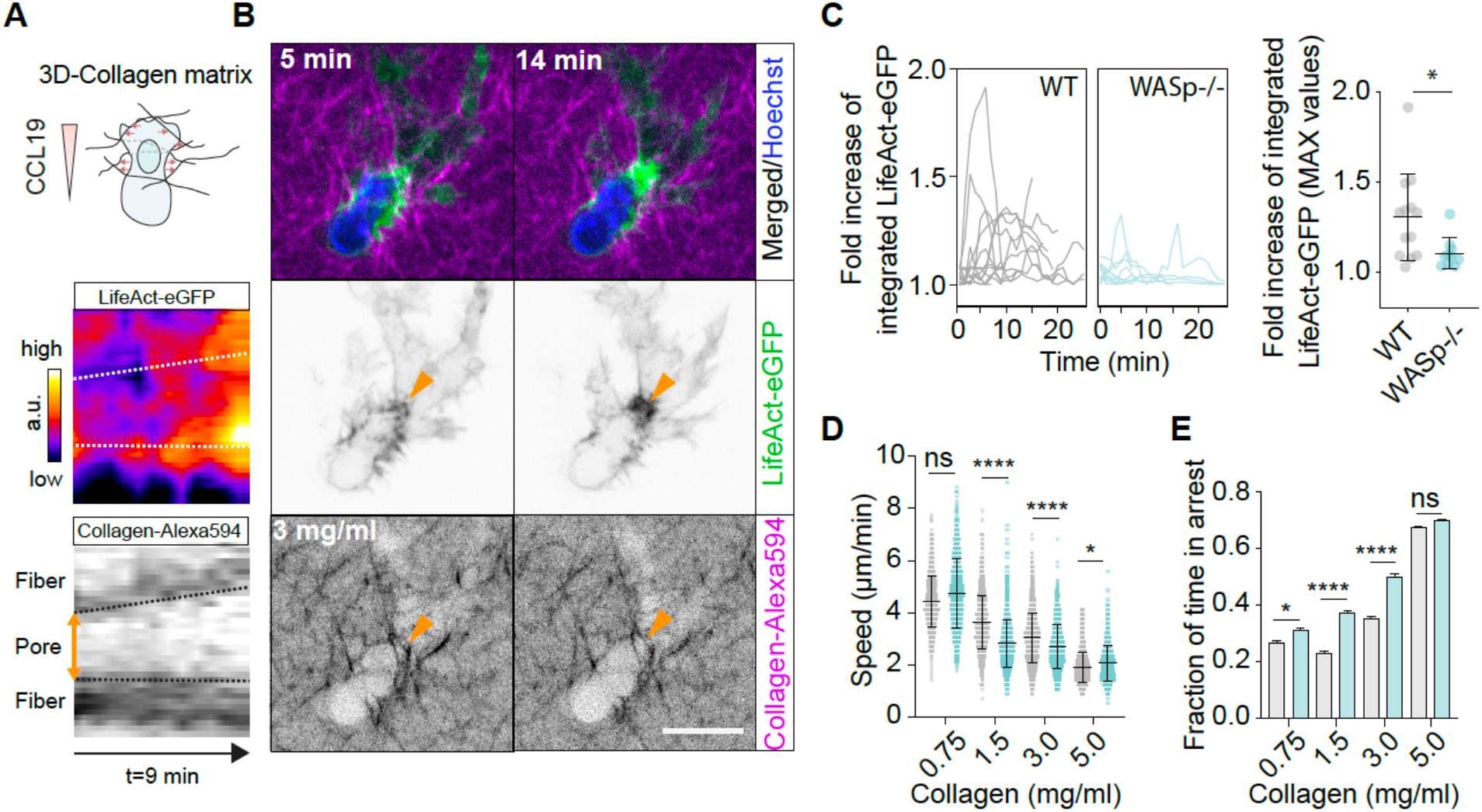
Deformation of collagen fibers is required for migration in fibrous environments. (**A**) Schematic of experimental setup. (**B**) Time-lapse spinning disc confocal microscopy reveals local deformation of obstructive collagen fibers (Alexa-594 labeled; 3.0 mg/ml) and increase of pore sizes for passage of the cell body (lower panel). Arrow heads indicate pore. Kymograph (left) shows dilation of the pore. Notably, fiber deformation was accompanied by local actin polymerization (middle panel). Upper panel shows merged images of Lifeact-eGFP and Collagen-Alexa-594. Scale bar=10 μm. (**C**) Left: Time-traces of LifeAct-eGFP signal at sites of obstructive fibers (normalized to intensity of the first frame; fold change of intensity is shown). Right: Each data point shows the local maximum of Lifeact-eGFP signal from one trace (fold change of intensity; first frame as reference). WT: n=12 cells; WASp−/−: n=10 cells; Mean±SD; Mann-Whitney test; *: p=0.0358. (**D**, **E**) Bulk analysis of DCs migrating in fibrous 3D collagen matrices with increasing densities. (D) shows mean speed of tracks pooled from 3 experiments. WT (0.75: n=694; 1.5: n=1181; 3.0: n=1551; 5.0: n= 981); WASp−/− (0.75: n=599; 1.5: n=1130; 3.0: n=1086; 5.0: n=853). Mean±SD; Kruskal-Wallis-test/Dunn’s multiple comparisons test; *: p=0.0221; ****: p<0.0001. (E) Fraction of time in arrest was extracted from tracks in (D); Mean+SEM; Kruskal-Wallis/Dunn’s multiple comparisons test; *: p=0.0470; ****: p<0.0001. ns: not significant. Also see Figure S5 and Movie S7.

### Orthogonal pushing drives T cell migration in crowded lymph nodes

WASp deficiency affects multiple hematopoietic lineages however, the dominant phenotype of WAS patients is a congenital immunodeficiency due to defective T-cell functions (Ochs, 2001). Cell migration is inevitably linked to activation of naïve T cells, as T cells need to constantly scan cell-packed secondary lymphoid tissues to search for cognate antigen on antigen presenting cells (Krummel et al., 2016). While WASp plays an established role in T cell development and immune synapse formation its role in interstitial T cell migration remained unclear (Thrasher and Burns, 2010). To test if the mechanism we established for DCs is relevant for lymphocytes, we purified naïve T cells from WASp−/− mice and confined them under agarose (10kPa). Locomotion was significantly impaired compared to WT controls and cells were frequently arrested under the load of the compressive overlay (**Figure 8A, Movie 6**). Similar to confined DCs, WT T cells formed highly dynamic actin foci traveling backward with the retrograde actin flow (**Figure 6B**). These actin foci were virtually absent in WASp−/− cells **(Figure 6B, Movie S6)**. Subcellular mechanical loading with both micropipette tips (**Figure S6A, Movie 8**) and nanometer sized ridges (**Figures 6C, S6B, Movie S8**) triggered the formation of WASp-dependent actin foci. Finally, cortical stiffness measured by submicron indentation with the tip of an AFM cantilever was significantly reduced in WASp−/− compared to WT cells (**Figures 6D, S6C**). Together, these data support a crucial role of WASp in the cortical mechanics of T cells. Next, we adoptively transferred labeled WT and WASp−/− naïve T cells into WT recipient mice **(Figure 6E)**. Since WASp−/− T cells show unimpaired homing to popliteal lymph nodes (Snapper et al., 2005), these experimental conditions allowed us to compare intranodal migration of both genotypes side-by-side **(Figure 6E, Movie S9)**. WASp−/− T cells were able to reach maximum speeds comparable to WT controls, indicating that WASp was not strictly required for locomotion of T-cells *in vivo* **(Figure S6D).**However, a significantly larger fraction of cells showed periods of minimal displacement (<2.5 μm in 1 min) **(Figure 6F)**, resulting in an overall reduced mean speed **(Figures 6G, S6E)**. These data confirm that WASp-dependent actin polymerization is of general relevance for 3D migration under compressive loads across both myeloid and lymphoid hematopoietic lineages and provide a mechanistic framework to defective cell migration observed in X-linked WAS (Bouma et al., 2009).

**Figure 6.**
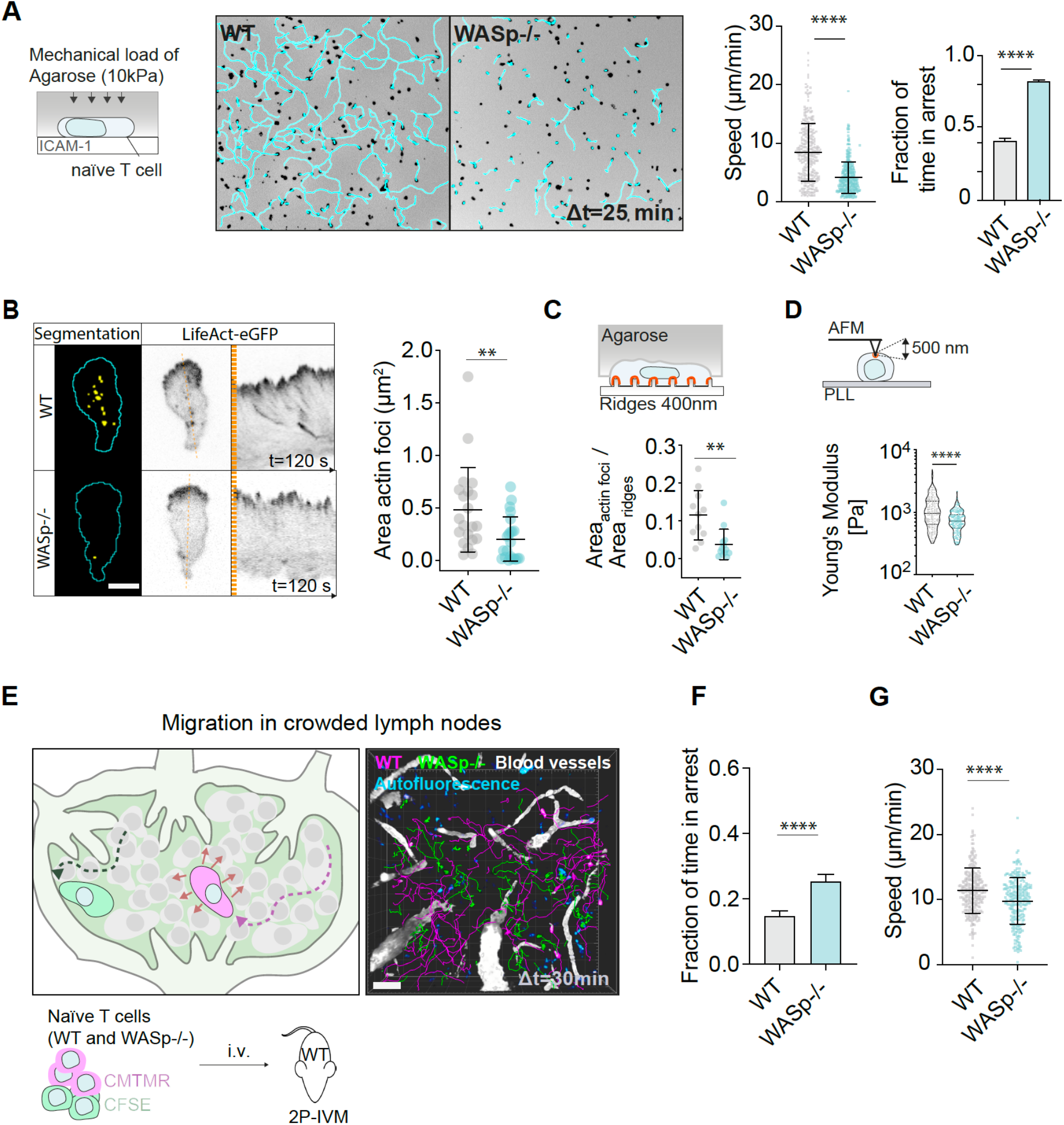
Orthogonal pushing drives T cell migration in crowded lymph nodes. Schematic of experimental setup. Epifluorescence images (CMTMR) from representative time series: Single cell tracks are overlaid in light-blue; scale bar=100 μm. Mean track speed was quantified. Each data point represents one track. WT: n=290, WASp−/−: n=335 pooled from 4 experiments; Mann-Whitney test; ****: p<0.0001. Fraction of time in arrest was extracted from tracks; Mean+SEM; Mann-Whitney test; ****: p<0.0001. (**B**) Actin foci were segmented from LifeAct-eGFP movies (spinning disc confocal microscopy) in Ilastik and area was quantified. Actin foci traveled backwards with the retrograde actin flow (kymographs); WT: n=21 cells, WASp−/−: n=22 cells; Mean±SD; Mann-Whitney test; **: p=0.0030; scale bar=5 μm. (**C**) Fraction of ridges covered by actin foci (segmented in Ilastik) (also see Figures S4B,S6B). WT: n=11; WASp−/−: n=13; Mean±SD; Mann-Whitney test; **: p=0.0015. (**D**) Schematic of an AFM experiment. Using a Hertz contact mechanics model, the elastic modulus was estimated by fitting the force indentation curves up to 500 nm. Each data point represents one measurement. WT: n=182 from 16 cells; WASp−/−: n=165 from 15 cells. For cellwise analysis see Figure S6C. Violin plot with Median±Quartiles; Mann-Whitney test; ****: p<0.0001. (**E**) Adoptive transfer of fluorescently labeled T cells (WT and WASp−/−) into wild-type recipient mice. After 24h, T cell migration in LN parenchyma was analyzed using intravital 2-photon microscopy. Representative 3D reconstruction of adoptively transferred T cells migrating in LN parenchyma (2-photon intravital microscopy). Scale bar=50 μm. (**F**) Fraction of time in arrest was extracted from tracks (see h); Mean+SEM; Mann-Whitney test; ****: p<0.0001. Mean track speed. Each data point represents one track. WT: n=262, WASp−/−: n=237 pooled from 5 experiments; Mann-Whitney test; ****: p<0.0001. Also see Figure S6 and Movies S8 and S9.

## DISCUSSION

First described in 1937, WAS is one of the most thoroughly investigated severe congenital immunodeficiencies (Ochs, 2001). WAS patients suffer from thromobocytopenia, eczema, and recurrent bacterial infections. Although underlying defects in cell-cell interactions and motility of the hematopoietic compartment have been attributed to the well-established function of WASp as one of the upstream activators of Arp2/3 nucleated actin polymerization, the precise cell biology behind WAS remained obscure (Graziano and Weiner, 2014; Machesky and Insall, 1998; Thrasher and Burns, 2010). We here show that WASp drives cortical actin polymerization in the third dimension and that it does so in a mechanosensitive manner. WASp dependent forces act orthogonal to the direction of cellular locomotion, and they are dispensable when the cell migrates in environments where the pore size is sufficiently large for the cell body to passage. WASp becomes rate-limiting for migration when the cell needs to pass through restrictive environments that require the cell to deform its viscoelastic surrounding. WASp assembles into dot-like structures that are embedded into the actin cortex and act as nucleation sites for spikes and ridges that then protrude against the external obstacle (**Figure S7)**.

This function of WASp contrasts with that of the Arp2/3-activating WAVE complex. The WAVE complex does not assemble in discrete spots but in traveling waves: as WAVE activity is self-amplifying and at the same time negatively regulated by polymerized actin, it travels and expands as a linear front of actin assembly (Graziano and Weiner, 2014). Such an “in-plane of the membrane” polymerization pattern only allows protrusion at the tip of a strictly flat lamellipodium, where the sheet of plasma membrane curves to form an envelope. Accordingly, the orientation of Arp2/3 branches in lamellipodia is exclusively horizontal (Svitkina, 2018). WASp lacks the self-organizing feature of the WAVE complex. It stays confined in dot-like assemblies, and its lateral movement seems restricted to passive co-migration within the surrounding cortex. This lack of self-organization might allow WASp activity to “remain on target” when the cell pushes against a target as delicate and small as a collagen fiber. When cells migrated on the nanotopographies, the precision with which the WASp dots located to the ridges was remarkable and suggested that WASp assembles in response to a highly localized signal. Importantly, this putative signal is independent of adhesive interactions, which is well in line with the amoeboid principle that can work in the absence of any cognate adhesive interaction with a substrate. Topographical changes of the plasma membrane as sensed e.g., by BAR domain-containing proteins are the most attractive candidates for such an alternative sensory function (Lou et al., 2019; Suetsugu and Gautreau, 2012).

How cells adapt to compressive load in 3D tissue environments is still incompletely understood. Recent studies proposed an evasion reflex in response to cellular compression: upon deformation of the nucleus cells increase their cortical contractility to move away and squeeze out from tight spaces or crowded tissue regions (Lomakin et al., 2020; Venturini et al., 2020). To reach their target sites, directionally migrating leukocytes, however, often need to enter and traverse tissues of high density and switch to an invasive, rather than an escape program is mandatory to remain on track. Formation of stiff actin spikes that push against the obstructing barrier may reflect the first, instantaneous response of a cell adopting an invasive mode of migration. Indeed, until to date, one of the best-established functions of WASp and its non-hematopoietic isoform N-WASP are invasive podosomes and invadosomes (Mooren et al., 2012; Murphy and Courtneidge, 2011). Although on the cellular scale, the WASp function we describe has little to do with an invasive process, the mechanistic aspect of the WASp-driven actin foci is similar to an extremely rudimentary and stalled podosome-driven invasion process. Podosomes are adhesive organelles, which are composed of a WASp and Arp2/3 containing protrusive core, surrounded by an adhesive ring-structure composed of integrins and their force coupling adaptors (van den Dries et al., 2019). A further defining feature of podosomes is their matrix-degrading activity, due to a podosome-targeted delivery of vesicles containing proteolytic enzymes of the matrix metalloproteinase family (Linder, 2007). Although in transcellular extravasating lymphocytes (Carman et al., 2007), immunological synapses (Kumari et al., 2015) and cells that undergo fusion (Sens et al., 2010), non-proteolytic podosome-like organelles were described, these structures were strictly coupled to (cell-cell) adhesions that presumably counter the pushing force of the protrusive actin-core. It seems plausible that in amoeboid migrating leukocytes, where integrins are either inactive or transmit horizontal traction forces (Hons et al., 2018) but never vertical adhesion forces, WASp-mediated protrusive foci are not locked in place by an adhesive ring and thus lack a maturation signal that turns them into degradative structures. Notably, work in invasive C elegans anchor cells showed that in WASp-dependent sites of basement membrane invasion, protrusive forces and proteolytic degradation synergize to drive barrier-penetration (Caceres et al., 2018). Upon inhibition of proteolysis, invasion can partially proceed in an entirely mechanically driven fashion (Kelley et al., 2019).

The protrusive actin foci we describe also share some features with early stages of clathrin-mediated endocytosis, where the positive curvature of the plasma membrane is associated with WASp-driven actin polymerization (Almeida-Souza et al., 2018). Actin polymerization on clathrin coated pits (CCP) is oriented normal to the cell surface, allowing it to push the CCP inward (Collins et al., 2011; Picco et al., 2015). This orientation is consistent with our findings that WASp plays a unique role in driving cortical actin polymerization in the third dimension. Of note, work in MDA-MB-231 cells showed that the endocytic machinery with its clathrin-based curved scaffolds can be diverted to wrap the plasma membrane around collagen fibers and to support mesenchymal cell migration (Elkhatib et al., 2017). Thus, the protrusive actin structures we describe share features with early stages of endocytosis as well as early stages of podosome formation. Together, our data support the general notion that WASp-driven actin polymerization mediates cellular mechanics acting orthogonal to the plasma membrane rendering cells mechanosensitive in 3D.

## ACKNOWLEDGEMENTS

We thank N. Darwish-Miranda, F. P. Assen, A. Kopf and A. Eichner for advice and help with experiments. We thank J. Renkawitz, E. Kiermaier, A. Juanes Garcia and M. Avellaneda for critical reading of the manuscript. We thank M. Driscoll for advice on fluorescent labelling of collagen gels. Further, we thank the Scientific Service Units of IST Austria for excellent technical support.

## AUTHOR CONTRIBUTIONS

Conceptualization, F.G., M.S; Methodology, F.G., P.R-R., I.V., M.H., J.A., J.M., V.Z., W.A.K.; Investigation, F.G., P.R-R., I.V., M.H.; Software, R.H.; Resources, A.L., Formal Analysis, F.G., R.H., P.R-R., M.R.; Writing – Original Draft, F.G. and M.S.; Writing – Editing, all authors; Visualization, F.G.; Supervision, F.G. and M.S.; Project Administration, F.G.; Funding Acquisition, F.G. and M.S.

## REFERENCES

Almeida-Souza, L., Frank, R.A.W., Garcia-Nafria, J., Colussi, A., Gunawardana, N., Johnson, C.M., Yu, M., Howard, G., Andrews, B., Vallis, Y., et al. (2018). A Flat BAR Protein Promotes Actin Polymerization at the Base of Clathrin-Coated Pits. Cell 174, 325–337 e314.

Barry, N.P., and Bretscher, M.S. (2010). Dictyostelium amoebae and neutrophils can swim. Proc Natl Acad Sci U S A 107, 11376–11380.

Benesch, S., Lommel, S., Steffen, A., Stradal, T.E., Scaplehorn, N., Way, M., Wehland, J., and Rottner, K. (2002). Phosphatidylinositol 4,5-biphosphate (PIP2)-induced vesicle movement depends on N-WASP and involves Nck, WIP, and Grb2. J Biol Chem 277, 37771–37776.

Berg, S., Kutra, D., Kroeger, T., Straehle, C.N., Kausler, B.X., Haubold, C., Schiegg, M., Ales, J., Beier, T., Rudy, M., et al. (2019). ilastik: interactive machine learning for (bio)image analysis. Nat Methods 16, 1226–1232.

Blumenthal, D., Chandra, V., Avery, L., and Burkhardt, J.K. (2020). Mouse T cell priming is enhanced by maturation-dependent stiffening of the dendritic cell cortex. eLife 9, e55995.

Bouma, G., Burns, S., and Thrasher, A.J. (2007). Impaired T-cell priming in vivo resulting from dysfunction of WASp-deficient dendritic cells. Blood 110, 4278–4284.

Bouma, G., Burns, S.O., and Thrasher, A.J. (2009). Wiskott-Aldrich Syndrome: Immunodeficiency resulting from defective cell migration and impaired immunostimulatory activation. Immunobiology 214, 778–790.

Bovellan, M., Romeo, Y., Biro, M., Boden, A., Chugh, P., Yonis, A., Vaghela, M., Fritzsche, M., Moulding, D., Thorogate, R., et al. (2014). Cellular control of cortical actin nucleation. Curr Biol 24, 1628–1635.

Caceres, R., Bojanala, N., Kelley, L.C., Dreier, J., Manzi, J., Di Federico, F., Chi, Q., Risler, T., Testa, I., Sherwood, D.R., et al. (2018). Forces drive basement membrane invasion in Caenorhabditis elegans. Proc Natl Acad Sci U S A 115, 11537–11542.

Carman, C.V., Sage, P.T., Sciuto, T.E., de la Fuente, M.A., Geha, R.S., Ochs, H.D., Dvorak, H.F., Dvorak, A.M., and Springer, T.A. (2007). Transcellular diapedesis is initiated by invasive podosomes. Immunity 26, 784–797.

Chugh, P., Clark, A.G., Smith, M.B., Cassani, D.A.D., Dierkes, K., Ragab, A., Roux, P.P., Charras, G., Salbreux, G., and Paluch, E.K. (2017). Actin cortex architecture regulates cell surface tension. Nat Cell Biol 19, 689–697.

Collins, A., Warrington, A., Taylor, K.A., and Svitkina, T. (2011). Structural organization of the actin cytoskeleton at sites of clathrin-mediated endocytosis. Curr Biol 21, 1167–1175.

de Noronha, S., Hardy, S., Sinclair, J., Blundell, M.P., Strid, J., Schulz, O., Zwirner, J., Jones, G.E., Katz, D.R., Kinnon, C., et al. (2005). Impaired dendritic-cell homing in vivo in the absence of Wiskott-Aldrich syndrome protein. Blood 105, 1590–1597.

Derivery, E., and Gautreau, A. (2010). Generation of branched actin networks: assembly and regulation of the N-WASP and WAVE molecular machines. Bioessays 32, 119–131.

Elkhatib, N., Bresteau, E., Baschieri, F., Rioja, A.L., van Niel, G., Vassilopoulos, S., and Montagnac, G. (2017). Tubular clathrin/AP-2 lattices pinch collagen fibers to support 3D cell migration. Science 356.

Friedl, P., and Wolf, K. (2010). Plasticity of cell migration: a multiscale tuning model. J Cell Biol 188, 11–19.

Fritz-Laylin, L.K., Lord, S.J., Kakley, M., and Mullins, R.D. (2018). Concise Language Promotes Clear Thinking about Cell Shape and Locomotion. Bioessays 40, e1700225.

Fritz-Laylin, L.K., Riel-Mehan, M., Chen, B.C., Lord, S.J., Goddard, T.D., Ferrin, T.E., Nicholson-Dykstra, S.M., Higgs, H., Johnson, G.T., Betzig, E., et al. (2017). Actin-based protrusions of migrating neutrophils are intrinsically lamellar and facilitate direction changes. Elife 6.

Gerard, A., Patino-Lopez, G., Beemiller, P., Nambiar, R., Ben-Aissa, K., Liu, Y., Totah, F.J., Tyska, M.J., Shaw, S., and Krummel, M.F. (2014). Detection of rare antigen-presenting cells through T cell-intrinsic meandering motility, mediated by Myo1g. Cell 158, 492–505.

Graziano, B.R., and Weiner, O.D. (2014). Self-organization of protrusions and polarity during eukaryotic chemotaxis. Curr Opin Cell Biol 30, 60–67.

Guimarães, C.F., Gasperini, L., Marques, A.P., and Reis, R.L. (2020). The stiffness of living tissues and its implications for tissue engineering. Nature Reviews Materials 5, 351–370.

Heinemann, F., Doschke, H., and Radmacher, M. (2011). Keratocyte lamellipodial protrusion is characterized by a concave force-velocity relation. Biophys J 100, 1420–1427.

Hons, M., Kopf, A., Hauschild, R., Leithner, A., Gaertner, F., Abe, J., Renkawitz, J., Stein, J.V., and Sixt, M. (2018). Chemokines and integrins independently tune actin flow and substrate friction during intranodal migration of T cells. Nat Immunol 19, 606–616.

Hu, J., Chen, S., Hu, W., Lu, S., and Long, M. (2019). Mechanical Point Loading Induces Cortex Stiffening and Actin Reorganization. Biophys J 117, 1405–1418.

Kelley, L.C., Chi, Q., Caceres, R., Hastie, E., Schindler, A.J., Jiang, Y., Matus, D.Q., Plastino, J., and Sherwood, D.R. (2019). Adaptive F-Actin Polymerization and Localized ATP Production Drive Basement Membrane Invasion in the Absence of MMPs. Dev Cell 48, 313–328 e318.

Krause, M., and Gautreau, A. (2014). Steering cell migration: lamellipodium dynamics and the regulation of directional persistence. Nature reviews Molecular cell biology 15, 577–590.

Krummel, M.F., Bartumeus, F., and Gerard, A. (2016). T cell migration, search strategies and mechanisms. Nat Rev Immunol 16, 193–201.

Kumari, S., Depoil, D., Martinelli, R., Judokusumo, E., Carmona, G., Gertler, F.B., Kam, L.C., Carman, C.V., Burkhardt, J.K., Irvine, D.J., et al. (2015). Actin foci facilitate activation of the phospholipase C-gamma in primary T lymphocytes via the WASP pathway. Elife 4.

Laevsky, G., and Knecht, D.A. (2003). Cross-linking of actin filaments by myosin II is a major contributor to cortical integrity and cell motility in restrictive environments. J Cell Sci 116, 3761–3770.

Lai, F.P., Szczodrak, M., Block, J., Faix, J., Breitsprecher, D., Mannherz, H.G., Stradal, T.E., Dunn, G.A., Small, J.V., and Rottner, K. (2008). Arp2/3 complex interactions and actin network turnover in lamellipodia. EMBO J 27, 982–992.

Lammermann, T., Bader, B.L., Monkley, S.J., Worbs, T., Wedlich-Soldner, R., Hirsch, K., Keller, M., Forster, R., Critchley, D.R., Fassler, R., et al. (2008). Rapid leukocyte migration by integrin-independent flowing and squeezing. Nature 453, 51–55.

Lang, N.R., Skodzek, K., Hurst, S., Mainka, A., Steinwachs, J., Schneider, J., Aifantis, K.E., and Fabry, B. (2015). Biphasic response of cell invasion to matrix stiffness in three-dimensional biopolymer networks. Acta Biomater 13, 61–67.

Leithner, A., Eichner, A., Muller, J., Reversat, A., Brown, M., Schwarz, J., Merrin, J., de Gorter, D.J., Schur, F., Bayerl, J., et al. (2016a). Diversified actin protrusions promote environmental exploration but are dispensable for locomotion of leukocytes. Nat Cell Biol 18, 1253–1259.

Leithner, A., Leithner, A., Merrin, J., Reversat, A., and Sixt, M. (2016b). Geometrically complex microfluidic devices for the study of cell migration. Protocol Exchange.

Leithner, A., Renkawitz, J., De Vries, I., Hauschild, R., Hacker, H., and Sixt, M. (2018). Fast and efficient genetic engineering of hematopoietic precursor cells for the study of dendritic cell migration. Eur J Immunol 48, 1074–1077.

Linder, S. (2007). The matrix corroded: podosomes and invadopodia in extracellular matrix degradation. Trends Cell Biol 17, 107–117.

Lomakin, A.J., Cattin, C.J., Cuvelier, D., Alraies, Z., Molina, M., Nader, G.P.F., Srivastava, N., Sáez, P.J., Garcia-Arcos, J.M., Zhitnyak, I.Y., et al. (2020). The nucleus acts as a ruler tailoring cell responses to spatial constraints. Science 370, eaba2894.

Lou, H.Y., Zhao, W., Li, X., Duan, L., Powers, A., Akamatsu, M., Santoro, F., McGuire, A.F., Cui, Y., Drubin, D.G., et al. (2019). Membrane curvature underlies actin reorganization in response to nanoscale surface topography. Proc Natl Acad Sci U S A 116, 23143–23151.

Machesky, L.M., and Insall, R.H. (1998). Scar1 and the related Wiskott–Aldrich syndrome protein, WASP, regulate the actin cytoskeleton through the Arp2/3 complex. Current Biology 8, 1347–1356.

Mogilner, A. (2006). On the edge: modeling protrusion. Curr Opin Cell Biol 18, 32–39.

Mooren, O.L., Galletta, B.J., and Cooper, J.A. (2012). Roles for actin assembly in endocytosis. Annu Rev Biochem 81, 661–686.

Murphy, D.A., and Courtneidge, S.A. (2011). The 'ins' and 'outs' of podosomes and invadopodia: characteristics, formation and function. Nat Rev Mol Cell Biol 12, 413–426.

Ochs, H.D. (2001). The Wiskott-Aldrich syndrome. Clinical Reviews in Allergy & Immunology 20, 61–86.

Paluch, E.K., Aspalter, I.M., and Sixt, M. (2016). Focal Adhesion-Independent Cell Migration. Annu Rev Cell Dev Biol 32, 469–490.

Pflicke, H., and Sixt, M. (2009). Preformed portals facilitate dendritic cell entry into afferent lymphatic vessels. J Exp Med 206, 2925–2935.

Picco, A., Mund, M., Ries, J., Nedelec, F., and Kaksonen, M. (2015). Visualizing the functional architecture of the endocytic machinery. Elife 4.

Pincus, Z., and Theriot, J.A. (2007). Comparison of quantitative methods for cell-shape analysis. Journal of Microscopy 227, 140–156.

Plotnikov, S.V., and Waterman, C.M. (2013). Guiding cell migration by tugging. Curr Opin Cell Biol 25, 619–626.

Pulecio, J., Tagliani, E., Scholer, A., Prete, F., Fetler, L., Burrone, O.R., and Benvenuti, F. (2008). Expression of Wiskott-Aldrich syndrome protein in dendritic cells regulates synapse formation and activation of naive CD8+ T cells. J Immunol 181, 1135–1142.

Redecke, V., Wu, R., Zhou, J., Finkelstein, D., Chaturvedi, V., High, A.A., and Hacker, H. (2013). Hematopoietic progenitor cell lines with myeloid and lymphoid potential. Nat Methods 10, 795–803.

Renkawitz, J., Kopf, A., Stopp, J., de Vries, I., Driscoll, M.K., Merrin, J., Hauschild, R., Welf, E.S., Danuser, G., Fiolka, R., et al. (2019). Nuclear positioning facilitates amoeboid migration along the path of least resistance. Nature 568, 546–550.

Renkawitz, J., Schumann, K., Weber, M., Lammermann, T., Pflicke, H., Piel, M., Polleux, J., Spatz, J.P., and Sixt, M. (2009). Adaptive force transmission in amoeboid cell migration. Nat Cell Biol 11, 1438–1443.

Reversat, A., Gaertner, F., Merrin, J., Stopp, J., Tasciyan, S., Aguilera, J., de Vries, I., Hauschild, R., Hons, M., Piel, M., et al. (2020). Cellular locomotion using environmental topography. Nature.

Riedl, J., Flynn, K.C., Raducanu, A., Gartner, F., Beck, G., Bosl, M., Bradke, F., Massberg, S., Aszodi, A., Sixt, M., et al. (2010). Lifeact mice for studying F-actin dynamics. Nat Methods 7, 168–169.

Rotty, J.D., Wu, C., and Bear, J.E. (2013). New insights into the regulation and cellular functions of the ARP2/3 complex. Nat Rev Mol Cell Biol 14, 7–12.

Schindelin, J., Arganda-Carreras, I., Frise, E., Kaynig, V., Longair, M., Pietzsch, T., Preibisch, S., Rueden, C., Saalfeld, S., Schmid, B., et al. (2012). Fiji: an open-source platform for biological-image analysis. Nat Methods 9, 676–682.

Sens, K.L., Zhang, S., Jin, P., Duan, R., Zhang, G., Luo, F., Parachini, L., and Chen, E.H. (2010). An invasive podosome-like structure promotes fusion pore formation during myoblast fusion. J Cell Biol 191, 1013–1027.

Sixt, M., and Lammermann, T. (2011). In vitro analysis of chemotactic leukocyte migration in 3D environments. Methods Mol Biol 769, 149–165.

Snapper, S.B., Meelu, P., Nguyen, D., Stockton, B.M., Bozza, P., Alt, F.W., Rosen, F.S., von Andrian, U.H., and Klein, C. (2005). WASP deficiency leads to global defects of directed leukocyte migration in vitro and in vivo. J Leukoc Biol 77, 993–998.

Suetsugu, S., and Gautreau, A. (2012). Synergistic BAR-NPF interactions in actin-driven membrane remodeling. Trends Cell Biol 22, 141–150.

Svitkina, T.M. (2018). Ultrastructure of the actin cytoskeleton. Curr Opin Cell Biol 54, 1–8.

Thiam, H.R., Vargas, P., Carpi, N., Crespo, C.L., Raab, M., Terriac, E., King, M.C., Jacobelli, J., Alberts, A.S., Stradal, T., et al. (2016). Perinuclear Arp2/3-driven actin polymerization enables nuclear deformation to facilitate cell migration through complex environments. Nat Commun 7, 10997.

Thrasher, A.J., and Burns, S.O. (2010). WASP: a key immunological multitasker. Nat Rev Immunol 10, 182–192.

Tinevez, J.Y., Perry, N., Schindelin, J., Hoopes, G.M., Reynolds, G.D., Laplantine, E., Bednarek, S.Y., Shorte, S.L., and Eliceiri, K.W. (2017). TrackMate: An open and extensible platform for single-particle tracking. Methods 115, 80–90.

Tsai, T.Y., Collins, S.R., Chan, C.K., Hadjitheodorou, A., Lam, P.Y., Lou, S.S., Yang, H.W., Jorgensen, J., Ellett, F., Irimia, D., et al. (2019). Efficient Front-Rear Coupling in Neutrophil Chemotaxis by Dynamic Myosin II Localization. Dev Cell 49, 189–205 e186.

van den Dries, K., Linder, S., Maridonneau-Parini, I., and Poincloux, R. (2019). Probing the mechanical landscape - new insights into podosome architecture and mechanics. J Cell Sci 132.

Vargas, P., Maiuri, P., Bretou, M., Saez, P.J., Pierobon, P., Maurin, M., Chabaud, M., Lankar, D., Obino, D., Terriac, E., et al. (2016). Innate control of actin nucleation determines two distinct migration behaviours in dendritic cells. Nat Cell Biol 18, 43–53.

Venturini, V., Pezzano, F., Català Castro, F., Häkkinen, H.-M., Jiménez-Delgado, S., Colomer-Rosell, M., Marro, M., Tolosa-Ramon, Q., Paz-López, S., Valverde, M.A., et al. (2020). The nucleus measures shape changes for cellular proprioception to control dynamic cell behavior. Science 370, eaba2644.

Walton, J. (1979). Lead asparate, an en bloc contrast stain particularly useful for ultrastructural enzymology. Journal of Histochemistry & Cytochemistry 27, 1337–1342.

Weigelin, B., Bakker, G.J., and Friedl, P. (2016). Third harmonic generation microscopy of cells and tissue organization. J Cell Sci 129, 245–255.

Wolf, K., Te Lindert, M., Krause, M., Alexander, S., Te Riet, J., Willis, A.L., Hoffman, R.M., Figdor, C.G., Weiss, S.J., and Friedl, P. (2013). Physical limits of cell migration: control by ECM space and nuclear deformation and tuning by proteolysis and traction force. J Cell Biol 201, 1069–1084.

Worbs, T., Hammerschmidt, S.I., and Forster, R. (2017). Dendritic cell migration in health and disease. Nat Rev Immunol 17, 30–48.

