## Supplemental Data for "Mechanosensitivity of amoeboid cells crawling in 3D"

**Methods**

Animals

Mouse strain: *C57BL/6* (Janvier); *WASp-/-* (B6.129S6-*Was*$^{tm1Sbs}$/J; No. 019458; The Jackson Laboratory); *LifeAct-eGFP* (Riedl et al., 2010)*; LifeAct-eGFPxWASp-/-; LifeAct-eGFPxHEM1-/-* (Leithner et al., 2016a). Mice were bred and maintained at the local animal facility in accordance with the IST Austria ethics commission and the Austrian law. Permission was granted by the Austrian Federal Ministry of Science, Research and Economy (identification code: BMWF-66.018/0005-II/3b/2012).

Cell culture

R10 medium, consisting of RPMI 1640, supplemented with 10% fetal calf serum (FCS), 2 mM L-glutamine, 100 U/ml penicillin, 100 μg/ml streptomycin, and 50 μM 2-mercaptoethanol (all Invitrogen), was used as basic medium for all cells unless stated otherwise. All cells were grown and maintained at 37 °C / 5% $CO_2$ unless noted otherwise. No cell lines used in this study were found in the database of commonly misidentified cell lines that is maintained by ICLAC and NCBI biosample. The cell lines were not authenticated. Cell lines tested negative for mycoplasma.

*Immortalized hematopoietic precursors*

The bone marrow of 6-12-wk-old mice was isolated and hematopoietic progenitor cell lines were generated by retroviral delivery of an estrogen-regulated form of HoxB8 (Leithner et al., 2018; Redecke et al., 2013).

*Differentiation of mature dendritic cells*

Dendritic cells (DCs) were differentiated from bone marrow (2.0 x $10^6$ BM-cells in 10 ml / dish) or transiently immortalized hematopoietic precursors (0.5 x $10^6$ precursor cells in 10 ml / dish) (Leithner et al., 2018; Redecke et al., 2013), both originating from 6-12-week-old, male or female C57BL/6J, LifeAct-eGFP, WASp-/-, LifeAct-eGFPxWASp-/- or LifeAct-eGFPxHem1-/- mice. Cells were differentiated in 9 ml R10, supplied with 1 ml in-house-generate granulocyte_macrophage colony-stimulating factor (GM-CSF) hybridoma supernatant. On day 3, 8 ml R10, supplied with 2 ml GM-CSF, was added. Half of the medium was replaced on day 6. On day 8, lipopolysaccharide (LPS) from *Escherichia coli* 0127:B8 (Sigma) was added to an end concentration of 200 ng/ml to mature DCs overnight.

*Plasmids and lentivirus production*

The following fusion constructs were used: eGFP-Abi1 (Lai et al., 2008), eGFP-WASp (Benesch et al., 2002). Fusion-protein-coding lentiviruses were produced in LX-293 HEK cell (Clontech) by co-transfection of the expression plasmids (pLenti6.3, Invitrogen) with pCMV-dR8.91 packaging- and pCMV-VSV-G envelope plasmids (a gift from B.Weinberg, MIT, USA, Addgene plasmid no. 8454).

*Transfection*

For transgene delivery, bone marrow-derived DCs were transfected with 4 μg DNA, according to manufacturer guidelines using the nucleofector kit for primary T cells (Amaxa; Lonza Group). Briefly, 5 × $10^6$ cells were resuspended in 100 µl reconstituted nucleofector solution and transferred to an electroporation cuvette, and a total amount of 4 µg plasmid DNA was added. Cells were transfected by using a protocol specifically designed for electroporating immature mouse DCs (program X-001). Transfected DCs were used one day after transfection and enriched by FACS for GFP-expressing cells. Immortalized hematopoietic precursors were spin-infected (1,500 g, 1 h) with lentiviruses in the presence of 8 μg/ml Polybrene. Following transduction, cells were selected for stable virus insertion using 10 µg/ml Blasticidin (pLenti6.3) for at least 1 wk.

*Purification of Naïve T-cells*

Peripheral LNs and spleens were harvested and homogenized with a 70 µm cell strainer. Untouched primary naïve T cells were isolated with an EasySep Mouse T cell Isolation Kit according to the manufacturer's protocol (STEMCELL Technologies, 19851 A).

Under-agarose migration assay

*Naïve T-cells*

T-cells were harvested 1 d before imaging and incubated overnight at 37 °C under 5% $CO_2$ in R10 medium consisting of RPMI 1640 supplemented with 10% FCS, 2 mM l-glutamine, 100 U/mL penicillin 100 µg/mL streptomycin, 1 mM sodium pyruvate, 100x nonessential amino acids and 50 µM 2-mercaptoethanol (all from Invitrogen). T cells were fluorescently labeled with 10 µM CMTMR (CellTracker Orange or 2 µM CFSE for 15 min at 37 °C.

Glass coverslips were overlaid with 4 µg/mL rmICAM-1/Fc (R&D, 796-IC). An 1% agarose block was formed by mixing (i) one part 2x HBSS buffer (Sigma), (ii) two parts RPMI (Invitrogen) supplemented with 20% BSA (instead of FCS) (Sigma) with 2x concentrations of all other supplements used in R10 medium (see above) and (iii) one part 4% high-molecular weight agarose (Biozym Gold Agarose, 850152) in water at 52 °C. CCL19 (20 ng/ml) (Peprotech, 250-27B) was added to soluble agarose before casting. Liquid agarose was subsequently poured into a dish, covering the coated coverslip. The agarose blocks were allowed to solidify at room temperature and were equilibrated first at 4 °C for 1 h and subsequently at 37 °C and 5% $CO_2$ for 30 min. T cells were injected under the agarose block with a micropipette and allowed to polarize for at least 30 min at 37 °C under 5% $CO_2$ before imaging. Epifluorescence movies were recorded using the same settings as described above. Images were taken every 30 s at 6 multi-positions with NIS Elements software (Nikon Instruments). Spinning disc microscopy was performed on an inverted spinning-disc confocal microscope (Andor) using a 100x/1.4 NA objective and a 488 nm laser line in a custom-built climate chamber (37°C under 5% $CO_2$). Time-lapse movies were recorded every two seconds.

*Mature Dendritic cells*

*Invasion under agarose.* Glass coverslips were washed with isopropanol, ethanol, and $dH_2O$; subsequently, plasma-cleaned (pdc-002 plasma cleaner, Harrick) and glued to a petri-dish with a 17mm hole. To obtain humid migration chambers, a 17 mm plastic ring was attached to a glass-bottom dish using paraffin (Paraplast X-tra; Sigma). Agarose blocks were formed by mixing (i) one part 2x HBSS buffer (Sigma), (ii) two parts RPMI (Invitrogen) supplemented with 20% FCS (Invitrogen) with 2x concentrations of all other supplements used in R10 medium (see above) and (iii) one part 4% UltraPure Agarose (Invitrogen) in water at 52 °C. Increasing agarose stiffnesses were achieved by mixing 2% agarose for 2.5 kPa, 4% for 10 kPa, and 6% for 15 kPa (Biozym Gold Agarose, 850152) (Hons et al., 2018). 500 µl liquid agarose was subsequently poured into a dish, covering the coverslip. The agarose blocks were allowed to solidify at room temperature for 5 min. After polymerization, two 2-mm holes (5 mm apart) were punched into the agarose pad followed by 30 min equilibration at 37°C, 5% $CO_2$. 2.5 µg/ml CCL19 (R&D Systems) was placed into one hole to generate a soluble chemokine gradient. 0.5 x $10^6$ mDCs were placed in the second hole opposite to the chemokine hole. Before the acquisition, dishes were incubated at least 2 h at 37°C, 5% $CO_2$ to allow invasion under the agarose (cells are now confined between the coverslip and the agarose). During the acquisition, dishes were held under physiological conditions at 37°C and 5% $CO_2$. Epifluorescence movies were recorded with an inverted wide-field Nikon Eclipse Ti-2 microscope in a humidified and heated chamber at 37 °C and 5% $CO_2$ (Ibidi Gas Mixer), equipped with a 20x/0.5 NA PH1 air objective, a Hamamatsu EMCCD C9100 camera and a Lumencor Spectra X light source (390 nm, 475 nm, 542/575 nm; Lumencor). Images were taken every 30 s or 3 min at 6 multi-positions with NIS Elements software (Nikon Instruments).

*Visualization of actin dynamics.* Plasma-treated (pdc-002 plasma cleaner, Harrick) glass coverslips were incubated with Poly-2-methyl-2-oxazoline (1 mg/ml for 1h at RT; PAcrAm™-g-(PMOXA); SuSoS Surface Technology) to generate an inert, non-adhesive coating. Agarose blocks were generated as described above. After polymerization (5 min at RT), a 2-mm hole was punched into the agarose pad followed by 30 min equilibration at 37°C, 5% $CO_2$. 2.5 µg/ml CCL19 (R&D Systems) was placed into the hole to generate a soluble chemokine gradient. The cell suspension was injected under the agarose opposite the chemokine hole to confine migrating DCs between the coverslip and the agarose. Before the acquisition, dishes were incubated at least 2 h at 37°C, 5% $CO_2$ to allow recovery and persistent migration of cells towards the chemokine source. TIRF microscopy was performed with a 60/1.46 NA oil objective, optovar 1x or 1.6x in a humidified and heated chamber at 37°C and 5% $CO_2$ using an inverted Axio Observer (Zeiss) microscope, a 488 nm laser and an Evolve EMCCD camera (Photometrics) controlled by VisiView software (Visitron Systems). Images were recorded every two seconds. Spinning disc microscopy was performed using the same settings as described for under agarose assays. To record actin spike formation z-stacks of 3 images (0.5 µm step size) were recorded every two seconds.

*Pharmacological inhibitors*

The following small molecule inhibitors were used to perturb actin dynamics. Inhibitors were mixed with the cell suspension and the agarose (before polymerization) using the indicated final concentration. The Arp2/3 Complex Inhibitor I, CK666 (100 µM; #SML0006; Sigma); the Arp2/3 Complex Inhibitor I, Inactive Control; CK689 (100 µM;

#US1182517; Merck Millipore); thee Formin FH2 Domain Inhibitor, SMIFH2 (40 µM; #344092; Merck Millipore).

Migration in microfabricated pillar forests and straight and constricted channels

Microfluidic devices with pillars were micro-fabricated with polydimethylsiloxane (PDMS) (Leithner et al., 2016b). Photomasks were designed using Coreldraw X18, printed on a chrome photomask (1 µm resolution; JD Photo data), followed by a spin coating step using SU-8 2005 (3,000 rpm, 30 s; Microchem) and a prebake of 3 min at 95°C. The wafer was then exposed to 100 $mJ/cm^2$ ultraviolet light on an EVG 610 mask aligner. After a postexposure bake of 3 min at 95°C, the wafer was developed in propylene glycol methyl ether acetate (PGMEA). A 1 h silanization with Trichloro(1H,1H,2H,2H-perfluorooctyl)silane was applied to the wafer. The devices were made with a 1:10 mixture of Sylgard 184 (Dow Corning), and air bubbles were removed with a desiccator. The PDMS was cured overnight at 85°C. Microdevices were attached to isopropanol/ethanol-cleaned coverslips and incubated for 1 h at 85°C after plasma cleaning (pdc-002 plasma cleaner, Harrick). Before the introduction of cells, devices were flushed and incubated with complete medium for at least 1 h. Figure S3D: Dimensions of the pillars were 5 x 30 µm (height x width). The spacing between pillars was 20 µm. Figuree 3I: 5 x 5 µm (height x width) and 1 µm, 2 µm or 3 µm spacing.

The dimensions of the straight channels are 5 µm width, 4 µm height and the dimensions of constrictions are 1.5 µm width, 4 µm height, 15 µm length. Brightfield movies of DCs migrating pillar mazes were acquired by time-lapse acquisition (time interval of 60 s) using inverted cell culture microscopes (DM IL Led, Leica Microsystems) equipped with cameras (ECO415MVGE, SVS-Vistek) and custom-built climate chambers (5% $CO_2$, 37°C, humidified).

Collagen migration assay

Custom-made migration chambers were assembled by using a plastic dish containing a 17-mm hole in the middle, which was covered by coverslips on each side of the hole (Sixt and Lammermann, 2011). 3D scaffolds consisting of 0.75 / 1.5 / 3 / 5 mg/ml bovine collagen I (PureCol, Nutragen, Fibricol; all AdvancedBioMatrix) were generated by mixing $1.5 \times 10^5$ cells in suspension (R10) with collagen I suspension buffered to physiological pH with Minimum Essential Medium and sodium bicarbonate in a 1:2 ratio. To allow polymerization of collagen fibers, gels were incubated 1 h at 37°C, 5% $CO_2$. Directional cell migration was induced by overlaying the polymerized gels with 0.63 µg/ml CCL19 (R&D Systems) diluted in R10. To prevent drying out of the gels, migration chambers were sealed with Paraplast X-tra (Sigma). Bright-field movies were acquired by time-lapse acquisition (time interval of 60 s) using inverted cell culture microscopes (DM IL Led, Leica Microsystems) equipped with cameras (ECO415MVGE, SVS-Vistek) and custom-built climate chambers (5% $CO_2$, 37°C, humidified).

To visualize collagen fibers using spinning disc microscopy, collagen was directly conjugated to Alexa Fluor 594 NHS Ester (Succinimidyl Ester, ThermoFisher). Collagen was added to SnakeSkin Dialysis Tubes, 10K MWCO, 16mm (ThermoFisher), and immersed in 100mM $NaHCO_3$ overnight at 4°C to allow polymerization. Alexa Fluor 594

NHS Ester (1.5mg/mL) was added to the polymerized collagen and incubated for 3h. To remove the unconjugated dye, the collagen mixture was placed in 0.2% acetic acid in deionized water for further dialysis overnight at 4°C.

Imaging of LifeAct-eGFP expressing DCs in Alexa-594-labeled collagen matrices was performed on an inverted spinning-disc confocal microscope (Andor Dragonfly 505) using a 60x/1.4 NA objective and a 488 nm / 561 laser line in a custom-built climate chamber (37°C under 5% $CO_2$). Z-stacks (1.5µm step size) of migrating cells in labeled collagen matrices were acquired using an Andor Zyla camera (4.2 Megapixel sCMOS) every 60 seconds for 20-25min.

Migration on nano-ridges

*Fabrication of coverslips with nano-ridges.* Clean glass coverslips were coated with a 50 nm reflective layer of chromium. An electron beam resist (AR-P 6200.13) was spin-coated on the chromium covered coverslip. The inverse pattern of ridges was written using the e-beam lithography tool and subsequently developed using AR600-546. Chromium was dry-etched in $Cl_2$-$O_2$ plasma inside an ICP (Inductively coupled plasma) chamber. The chamber was pumped to a pressure of 10 mTorr with a gas flow of 26 sccm for $Cl_2$ and 4 sccm for $O_2$. The forward power and ICP power were 20 W and 400 W, respectively. Glass was subsequently etched using a $SF_6$-Ar plasma inside an ICP chamber, pumped to a pressure of 10mTorr with a gas flow of 100 sccm for $SF_6$ and 67 sccm for Ar. The forward power and ICP power were 100 W and 1500 W, respectively. After dry-etching, the remaining chromium layer was wet etched at room temperature using Chromium etchant. The height of the ridges is determined by the etching time of the glass surface, on average the etching rate of the glass is around 118.5 nm/min. After the wet etching step, the height of the ridges is verified using an atomic force microscope (NX10 from Park Systems) in non-contact mode.

*Under agarose assay on nano-ridges.* Glass coverslips with ridges were rinsed with isopropanol, ethanol and $dH_2O$, air-dried and plasma cleaned (pdc-002 plasma cleaner, Harrick). They were then incubated with Poly-2-methyl-2-oxazoline (1 mg/ml for 1h at RT; PAcrAm™-g-(PMOXA); SuSoS Surface Technology) or PLL-PEG (1 mg/ml for 1h at RT; PLL-PEG; SuSoS Surface Technology) to generate an inert, non-adhesive coating. Agarose blocks (1%) were generated as described above and matured DCs or purified naïve t-cells were injected under the agarose using a micropipette. Spinning-disc confocal microscopy and TIRF microscopy were performed as described above.

Micropipette indentation assay

Glass bottom dishes (50 mm dish diameter, 14 mm glass diameter, glass coverslips No. 1, Mattek) were plasma cleaned (pdc-002 plasma cleaner, Harrick) and coated with 1x poly-L-lysine (P8920, Merck) in $dH_2O$ for 10 min. Dishes were washed twice with $dH_2O$ and then dried for at least 4h at room temperature. Cells in R10 (mDCs or t-cells expressing LifeAct-eGFP) were incubated for 15 min at 37 °C and dishes were carefully washed once with R10 containing HEPES (10mM; Sigma) to remove floating cells. Dishes were immediately mounted on an inverted spinning-disc confocal microscope (Andor) equipped with a micromanipulator (Eppendorf) and maintained at 37°C in a custom-built climate chamber. Micropipettes (blunt; inner diameter 4 µm; bent angle 30°) (BioMedical Instruments) were centrally positioned over the cell and carefully lowered to indent the cell body. Movies were recorded using a 100x/1.4 NA objective and a 488 nm laser line. Z-stacks of 3 image (0.5 µm step size) were recorded every two seconds.

Electron Microscopy, ultrastructural analysis

*Under-agarose assay, fixation and light microscopy examination.* Under-agarose assays were performed as described above with minor modifications. Removable culture inserts (Cat.No: 81176; Ibidi) were attached to plasma-cleaned (pdc-002 plasma cleaner, Harrick) glass coverslips or tailored Aclar foil (Science Services) (used for serial sectioning). Culture inserts were then filled with 200 µl of agarose-mix and experiments were performed as described above. Dendritic cells were allowed to migrate under the agarose for > 2h. Samples were subsequently fixed in cytoskeleton buffer (10 mM MES buffer, 150 mM NaCl, 5 mM EGTA, 5 mM glucose and 5 mM $MgCl_2$; pH=6.1) containing 2% PFA; 2.5% glutaraldehyde (EMS); 0.01% Triton-X 100 (Sigma); phalloidin-Alexa488 (1:40; Invitrogen) for 30 min at 37°C. After fixation, agarose pads and removable culture inserts were carefully removed using a coverslip tweezer. Z-Stacks of palloidin-Alexa488 stained cells were performed on an inverted spinning-disc confocal microscope (Andor) using a 100x/1.4 NA objective and a 488 nm laser line. For CLEM (correlative light and electron microscopy) epifluorescence images of phalloidin-Alexa488 stained cells were recorded with an inverted wide-field Nikon Eclipse Ti-2 microscope equipped with a 20x/0.5 NA PH1 air objective, a Hamamatsu EMCCD C9100 camera and a Lumencor

Spectra X light source (475 nm Lumencor). Images were taken at multi-positions with NIS Elements software (Nikon Instruments). Fluorescent and SEM images (see below) were manually aligned using FIJI.

*Scanning Electron Microscopy, surface analysis.* Samples were dehydrated in a graded ethanol series at RT. For further chemical drying, the samples were first incubated in a 1:1 mixture of 100% ethanol and hexamethyldisilazane (HMDS) for 30min at RT and then transferred to 100% HMDS for 1h at RT. Access HMDS was removed with a pipette followed by overnight evaporation at RT. The cover slips with the dried cells were coated with platinum to a thickness of 5nm using an EM ACE600 coating device (Leica Microsystems). The samples were observed with a FE-SEM Merlin compact VP scanning electron microscope (Zeiss) at 5kV using a secondary electron detector.

*Contrast enhancement and resin embedding.* Samples on Aclar foil were post-fixed with 1% glutaraldehyde (GA) (EMS) in phosphate buffer (PB; 0.1M, pH 7.4) for 10 min at RT. After a brief wash with PB, samples were contrast-enhanced with 0.5% tannic acid (TA; Sigma) in PB (w/v) for 45 min at 4 °C in the dark. The solution was replaced with freshly prepared 0.5% TA in PB and samples incubated for another 45 min at 4 °C in the dark. After a wash with PB, samples were incubated in 1% aqueous osmium tetroxide (w/v; EMS) for 30 min at 4 °C in the dark. Then, they were washed with Milli-Q water and incubated in 1% aqueous uranyl-acetate (w/v; AL-Labortechnik) overnight at 4 °C in the dark. After wash with Milli-Q water, samples were contrast-enhanced further according to en bloc Walton's lead aspartate staining (Walton, 1979). Samples were incubated in lead aspartate solution for 30 min at 60 °C. After a wash with Milli-Q water, samples were dehydrated in graded ethanol (10%, 20%, 50%, 70%, 90%, 96% and 100%) for 5 min each at 4 °C. They were then placed in propylene oxide puriss. p.a. (Sigma) twice for 10 min at 4 °C and embedded in Durcupan™ ACM epoxy resin (Sigma). To that, hard grade Durcupan™ was formulated by weight as follows: component A 11.4 g, component B 10 g, component C 0.3 g, component D 0.1 g. Samples were consecutively infiltrated in mixtures of 3:1 propylene oxide/Durcupan™, 1:1 propylene oxide/Durcupan™ and 1:3 propylene oxide/Durcupan™ for 1.5 h each at 4 °C. Then they were infiltrated in mere Durcupan™ overnight at RT. The Aclar foil was removed from the resin block with a razor blade under a stereo microscope (Nikon SMZ 800). Trimmed samples were placed on PELCO® Cavity Embedding Molds (Ted Pella Inc.), cavities filled with freshly prepared Durcupan™ and resin cured at 60 °C in an oven over two days.

*Serial sectioning and scanning.* Embedded samples were trimmed with an Ultratrim diamond knife (Diatome) to a rectangle (70µm x ~1mm) with a slanted side for orientation using an EM UC7 ultramicrotome (Leica Microsystems). Prior to serial sectioning, a 25 x 25mm silicon wafer was plasma treated using an ELMO glow discharge cleaning system (Agar Scientific) for increasing the hydrophilicity of the wafer. Serial sections were cut at a thickness of 80 nm using a 4mm Leica AT-4 35° diamond knife (Diatome). Section ribbons were collected onto a plasma treated wafer using the water drain device of the knife. Then the wafer with the serial sections was coated with carbon to a thickness of 5nm using an EM ACE600 coating device (Leica Microsystems) to ensure conductivity. The serial sections were observed under a FE-SEM Merlin compact VP scanning electron microscope (Zeiss) equipped with the Atlas 5.3.2.9 Array Tomography software (Zeiss). The images were acquired using a backscattered (5nm pixel size) and a secondary electron detector (7nm pixel size) at 5kV.

Atomic force microscopy

Glass bottom dishes (30 mm dish diameter, 14mm glass diameter, glass coverslip No. 1, Mattek) were plasma cleaned (pdc-002 plasma cleaner, Harrick) and coated with 1x poly-L-lysine (P8920, Merck) in $dH_2O$ for 10min. Dishes were washed twice with $dH_2O$ and then dried for at least 4h at RT. Cells in R10 (mDCs or t-cells expressing LifeAct-eGFP) were incubated for 15 min at 37°C and dishes were carefully washed once with R10 containing HEPES (10mM; Sigma) to remove floating cells. Dishes were immediately mounted on the atomic force microscope equipped with a climate chamber (37°C). AFM nanoindentation was performed on a Nanowizard4 AFM microscope from JPK Instruments (Bruker) interfaced to an inverted optical microscope (IX81, Olympus). We used cantilevers with spring constants of 0.1 N $m^{-1}$ and a tip radius of 10 nm (qp-BioAC, Nanosensors). Cantilever actual spring constants were determined using the thermal noise method implemented in the JPK software. Using brightfield microscopy, we positioned the tip of the cantilever over the central region above the cell body and performed 5-10 indentation measurements. Force-distance curves were acquired with an approach speed of 2 µm $s^{-1}$ until reaching the maximum set force of 10 nN. Measurements were restricted to indentation depths of 500 nm for T-cells and 1000 nm for DCs (<10% of the height of the cell) to minimize the contribution of the substrate, and to maximize the contribution of the cortex stiffness to cantilever deflection. Elastic moduli (young’s modulus) were determined from force-distance curves with the Hertz model as implemented in the JPK analysis software (parabolic model).

<u>Dendritic cell lymph node homing assay</u>

*Fluorescence labeling of dendritic cells.* Bone marrow-derived dendritic cells were differentiated and matured as described above. Mature DCs were harvested, counted and adjusted to 1x10$^{7}$ cells/ml at RT 1xPBS. For labeling, Tetra-Methylrhodamine (TAMRA; TAMRA, SE; 5-(and-6)-Carboxytetramethylrhodamine, Succinimidyl Ester (5(6)-TAMRA, SE; Invitrogen) and Oregon Green® (CellTrace™ Oregon Green® 488 carboxylic acid diacetate, succinimidyl ester carboxy-DFFDA, SE; Invitrogen) was added to a final concentration of 10 µM TAMRA or 3 µM Oregon Green®, respectively. After a 10 min incubation fresh R10 medium was added to the cell suspension to stop the reaction, and the cells were pelleted. Subsequently, cells were resuspended in pre-warmed (37°C) R10 medium and incubated for another 30 minutes at 37°C for esterification. Finally, cells were washed twice with 1xPBS and subsequently used for experiments. WT and WASp-/- were differently labeled, mixed at 1:1 ratio, and adjusted to a final concentration of 4x10$^{7}$ cells/ml in 1xPBS. 25 µl (= 1x10$^{6}$ cells) were injected into the mouse hind footpads. Draining popliteal lymph nodes (LNs) were harvested after 24 hours and transferred to a polystyrene FACS tube (Falcon BD) containing 0.5 ml complete DMEM (2.5% FCS (Gibco), 10 mM HEPES (Sigma-Aldrich), 5% penicillin/ streptomycin (Gibco) and 5 % glutamine (Gibco)) for subsequent isolation of the DCs for flow cytometry analysis (one LN per tube). LNs were then opened and cut into pieces in the tube using sterile scissors. Afterwards, LN fragments were incubated in digestion buffer (complete DMEM, 3mM $CaCl_2$ (Sigma-Aldrich), 0.5 mg/ml collagenase D (Roche), 40 µg/ml DNAse I (Roche) EDTA 0.5 M, pH 7.2: Ethylenediaminetetraacetic acid (EDTA, Sigma-Aldrich)) for 30 min at 37 °C in a water bath and solution was thoroughly pipetted every 10 minutes using a 1 ml pipette to further disrupt the fragments. The enzymatic reaction was stopped after 30 minutes by the addition of 10 mM EDTA. The cell solution was again thoroughly pipetted to disrupt remaining LN fragments and subsequently flushed through a cell strainer into a fresh FACS tube (tube with cell strainer cap, BD Falcon). After washing with FACS-buffer, cells were directly stained with labeled primary antibodies (CD11c (Antigen: CD11c; conjugated: APC; eBioscience; Cat-No 17-0114-82; Clone N418) and mouse MHC II (I-A/I-E) (Antigen: MHC II; conjugated: eFluor450; eBioscience; Cat-No 48-5321-82; Clone M5/114.15.2)), resuspended in an appropriate volume of FACS buffer and used for FACS analysis. Flow-cytometric analysis was performed on FACS Canto II (Becton Dickinson) or FACS Aria III (Becton Dickinson) using FACS DIVA software (Becton Dickinson) for acquisition and FlowJo (Treestar) for analysis.

Intravital two-photon microscopy of popliteal lymph nodes

Freshly purified T cells were fluorescently labeled with 20 µM CMTMR (CellTracker Orange or 5 µM CFSE for 15 min at 37 °C. After being washed, labeled T cells were i.v. injected retro-orbitally into sex-matched 5- to 10-week-old WT C57BL/6 recipient mice and were allowed to home to lymph nodes for at least 18 h. Recipient mice were anesthetized by intraperitoneal injections of ketamine (50 mg $kg^{-1}$), xylazine (10 mg $kg^{-1}$) and acepromazine (4 mg $kg^{-1}$). The right popliteal lymph node was prepared micro-surgically for intravital microscopy and positioned on a custom-built microscope stage. Care was taken to spare blood vessels and afferent lymph vessels. The prepared LN was submerged in normal saline and covered with a glass coverslip. A thermocouple was placed next to the LN to monitor local temperature, which was maintained at 37°C. To label blood vessels, 50 μl of Evans blue (1mg/ml) were i.v. injected retro-orbitally before imaging.Two-photon microscopy of right popliteal LNs was performed with a Trimscope II multi-photon imaging platform (LaVision Biotech) on an upright Olympus stand. Images were acquired using a Plan-Apochromat 20x/1.0 NA objective (Carl Zeiss Microscopy) with $H_2O$ as an immersion medium. A Chameleon Ti:Sapphire laser (MaiTai) was tuned to 840 nm, and an optical parametric oscillator (OPO) tuned to 1100 nm for simultaneous excitation of CFSE, CMTMR, and Evans blue. Fluorescent signals were collected using four external/non-descanned photomultipliers (PMTs) (3 Hamamatsu H7422-40 GaAsP High Sensitivity PMTs and 1 Hamamatsu H7422-50 GaAsP High Sensitivity red-extended PMT). For four-dimensional analysis of cell migration, z-stacks with 11–25 slices (spacing 4µm) of 250–300 x 250–300 µm *x–y* sections were acquired every 20 s for 30–45 min.

Image analysis

FIJI imaging processing software (https://fiji.sc/) was used for basic image and video microscopy analysis (Schindelin et al., 2012).

*Cell tracking*

*3D analysis of T-cell migration in LNs.* Imaging sequences of image stacks were transformed into volume-rendered four-dimensional movies using Imaris software (v9; Bitplane), which was also used for semiautomated tracking of cell motility in three dimensions. The average track velocity and instantaneous velocities were calculated from the x, y, and z coordinates of cell centroids.

*2D analysis of migration assays (under agarose; pillar forests; collagen assays)*

The average track velocity and instantaneous velocities were calculated from the x, y-coordinates of the nucleus (under agarose assays of dendritic cells) or the cell centroid (under agarose assay of T-cells / dendritic cells migrating in pillar forests or in collagen gels) tracked in TrackMate (Tinevez et al., 2017) (https://imagej.net/TrackMate).

*Fraction of time in arrest*. T-cells were classified as being in arrest when the cell centroid remains confined within a radius of 2.5 μm during an interval of 1 min. Dendritic cells were classified as being in arrest when the cell nucleus remains confined within a radius of 2.5 μm during an interval of 3 min. „Fraction of time in arrest“ was calculated by ratio of the time spent in arrest to the total time.

*Segmentation of actin foci*

To quantify the area and dynamics of actin foci we first segmented actin foci from LifeAct-eGFP expressing cells with interactive machine-learning using Ilastik (Berg et al., 2019). Training was performed on WT cells and the same trained workflows were then applied to WASp-/- cells. To measure the dynamic changes of actin foci from frame to frame, we calculated the Jaccard similarity coefficient, defined by the overlap of segmented actin foci of one frame with the segmentation of the previous frame divided by the area of the union of both. Hence, highly persistent actin foci would lead to a Jaccard coefficient of 1.

*Shape Analysis*

Dendritic cell bodies were segmented from fluorescent images (LifeAct-eGFP) by thresholding and convertion into binary images using FIJI (Schindelin et al., 2012). Polygonal outlines were extracted from segmentation masks and sampled at evenly spaced 200 points. To ensure that all polygons were orientated equally, an algorithm based on Procrustes analysis was used to rotate and translate the polygons until corresponding points were optimally aligned. We used the nucleus as a characteristic landmark defining the cell body of a migrating dendritic cell. Nuclei were extracted from fluorescent images (Hoechst) in FIJI and converted to another set of binary images. This additional landmark was used in the Procrustes procedure and improved the alignment. Finally, the alignment was manually verified. The following cellular characteristics were measured (1) aspect ratio = long axis/short axis, (2) the average diameter of the contour along the central axis, (3) length of the central axis and (4) normalized polygon curvature at the cell front was measured as an approximation of leading-edge roughness (The average of the absolute values of the point wise curvature of the contour is computed over a specified range, and multiplied by the contour length over the same range. Absolute values must be used because otherwise positive and negative curvatures would cancel out; the sum is multiplied by the arc length to make the measurement scaleinvariant.) All algorithms are implemented in the ’’Celltool’’ software package (Pincus and Theriot, 2007).

*Cross-correlation analysis and bootstrapping statistics*

To compare the temporal dynamics of two parameters (speed, aspect ratio, fluorescence of actin foci, nuclear area) and test for the statistical significance of the temporal offset we used cross-correlation analysis with a custom-written MATLAB script. Bootstrapping statistics was performed as described in (Tsai et al., 2019). Bootstrapping was used to obtain the 95% confidence interval of each cross-correlation value. Time traces of x and y were randomly permuted, and the same cross-correlation analysis to obtain the maximal correlation with all possible temporal offsets was performed. This process was then repeated 2000 times to obtain a distribution of maximal correlation value. The 95% confidence interval indicated a correlation value better than 95% of the maximal correlation values one can obtain with a pair of randomly permuted x and y.

*PIV of collagen matrices*

The displacement vectors of collagen fibers deformed by migrating DCs were calculated with the software Davis 8 (LaVision) applying Particle Image Velocimetry (PIV) on spinning-disc confocal images. Further post-processing was carried out using a custom-written Python script for extracting the maximum of deformation from frame-to-frame. Considered vectors were limited to the vicinity of the cell boundary corresponding to twice the cell area, which initially was determined by the LifeAct-eGFP expression.

Statistical analysis

Figures represent pooled data from independent biological replicates as indicated in the figure legends. Appropriate control experiments were performed for each biological replicate. All replicates were validated independently and pooled only when showing the same trend. Statistical analysis was conducted using Prism7 (GraphPad). D'Agostino Pearson omnibus K2 test was used to test for Gaussian or non-Gaussian data distribution, respectively. When data was normal distributed paired t-test was used. When data was non-normal distributed Kruskal-Wallis with Dunn's test or two-tailed Mann-Whitney test was used. Statistical tests used for individual experiments are indicated in the figure legends.

Data and software availability

All data is available in the main text or the supplementary materials. Computational analysis performed in this paper was performed using custom-written programs in Matlab (Mathworks) and Python. The analysis code will be made available upon request.

**Figure S1**

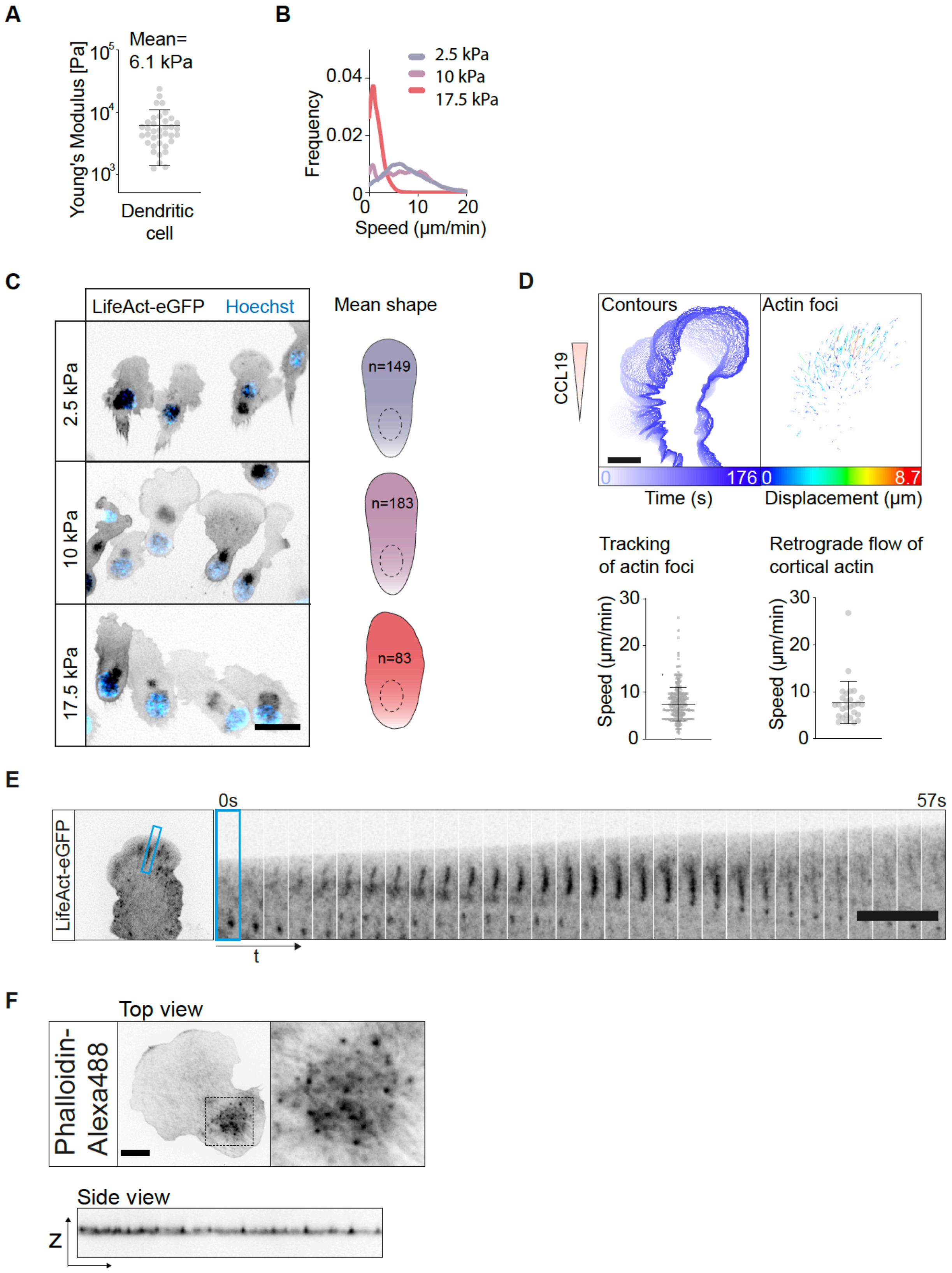
A
Mean=
6.1 kPa
Young's Modulus [Pa]
Dendritic
cell
B
2.5 kPa
10 kPa
17.5 kPa
Frequency
Speed (µm/min)
C
LifeAct-eGFP
Hoechst
Mean shape
2.5 kPa
10 kPa
17.5 kPa
n=149
n=183
n=83
D
Contours
Actin foci
CCL19
Time (s)
Displacement (µm)
Tracking
of actin foci
Retrograde flow of
cortical actin
Speed (µm/min)
E
LifeAct-eGFP
0s
57s
t
F
Top view
Phalloidin-
Alexa488
Side view
Z

**Figure S1. Related to Figure 1. Actin spikes form in response to mechanical loads of restrictive environments.** (**A**) Atomic force microscopy of mature DCs. Each data point represents one cell; n=37. (**B**) Frequency distribution of instantaneous cell speeds (smoothed histogram) under the load of agarose (also see Figure 1C). (**C**) Representative epifluorescence micrographs of DCs migrating under agarose of different stiffness; scale bar=30 μm. Cell contours were extracted from LifeAct-eGFP signal and mean cell shape calculated from all contours is shown (right panel). (**D**) Contours and actin foci (segmented from spinning disc confocal movies (LifeAct-eGFP) in Ilastik and tracked with TrackMate) of a representative cell; $\Delta t$=176 s; also see Movie S1). Color-code shows displacement of actin foci in μm. Scale bar=10 μm. Tracking of actin foci (n=481 tracks pooled from 2 cells) revealed mean speed comparable to the mean speed of bulk cortical actin flows as analyzed in kymographs (n=28 cells); Mean±SD. (**E**) Spinning disc confocal image sequence showing elongation of actin foci into stripes. Scale bar=10 μm. (**F**) Confocal z-scan of fixed cell stained with phalloidin-Alexa488; scale bar=10 μm.

### Figure S2

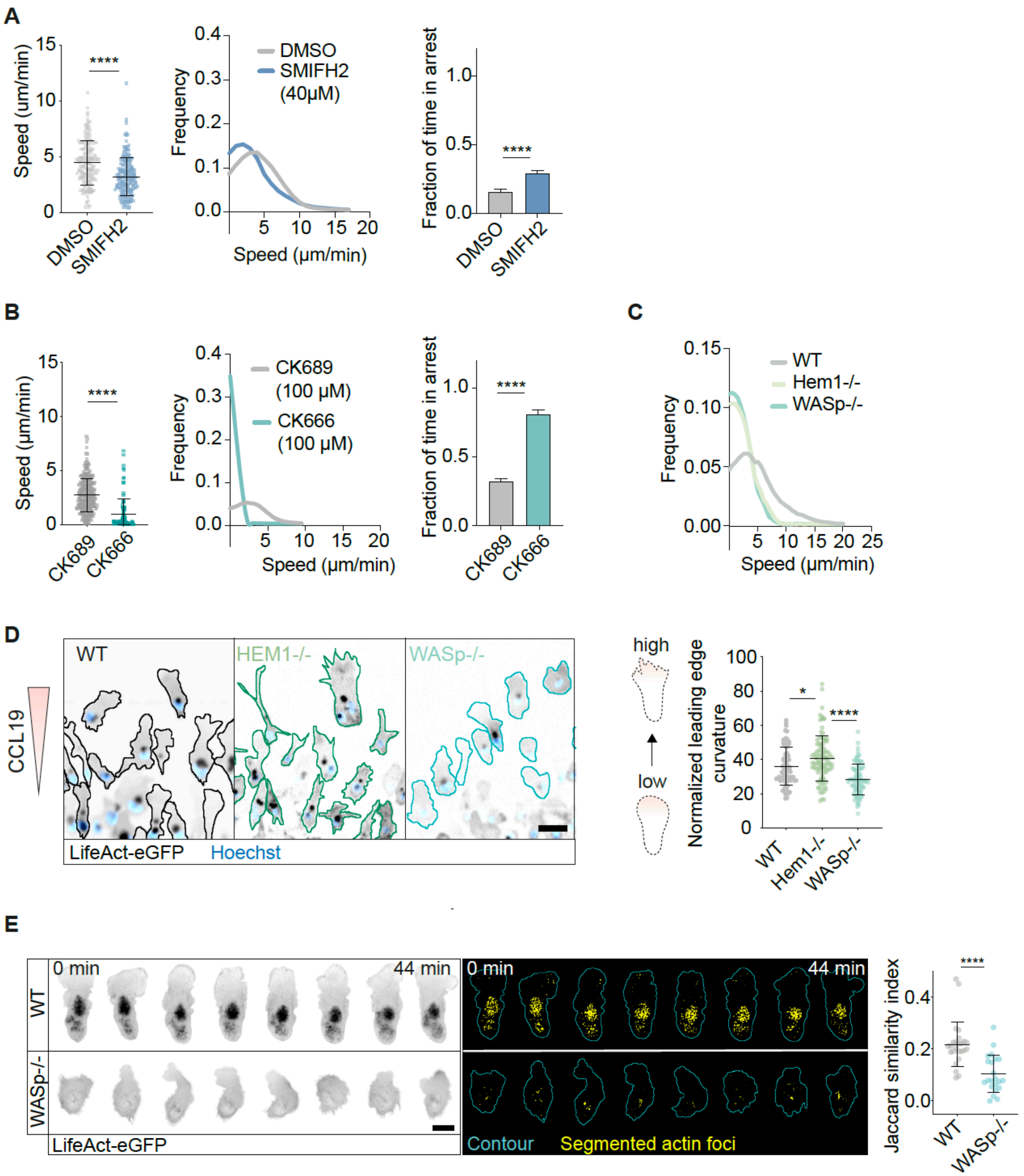


**Figure S2. Related to Figure 2. WASp-driven actin spikes polymerize orthogonal to WAVE-driven lamellipodia. (A, B)** WT DCs (Hoxb8-derived) were treated with small molecule inhibitors to block actin polymerization: Formin-inhibitor SMIFH2 (40 μM) and

Arp2/3-inhibitor (100 μM). Mean track speed of DCs migrating under agarose (10kPa). Each data point represents one track; Mean±SD. (**A**) DMSO: n=168; SMIFH2: n=193 pooled from 3 experiments; Mann Whitney test; ****: p<0.0001. (**B**) CK689 (inactive control): n=289 pooled from 4 experiments; CK666: n=105 pooled from 6 experiments; Mann Whitney test; ****: p<0.0001. Middle panels: Frequency distribution of instantaneous cell speeds (smoothed histogram). Right panels: Fraction of time in arrest was extracted from tracks; Mean+SEM; Mann Whitney test and Kruskal-Wallis/Dunn's multiple comparisons test; ****: p<0.0001. (**C**) Frequency distribution of instantaneous cell speeds (smoothed histogram). (**D**) Representative epifluorescence micrographs of DCs migrating under (10kPa); scale bar=50 μm. Cell contours were extracted from LifeAct-eGFP signal; WT: n=75; Hem1-/-: n=100; WASp-/-: n=84. Contours were further analyzed for leading edge curvature; Kruskal-Wallis/Dunn's multiple comparisons test; *: p=0.0466, ****: p<0.0001. (**E**) Time series of epifluorescence (LifeAct-eGFP) and segmented actin foci in WT and WASp-/- DCs (related to Figures 2 F,G). Cell contours were stabilized and motion of actin foci is shown in the cell-frame of reference. Scale bar=20 μm. Jaccard similarity index shows frame-to-frame overlap of segmented actin foci (mean of cell). WT: n=25 cells; WASp-/-: n=23 cells. Mean±SD; Mann-Whitney; ****: p<0.0001; ns=not significant.

**Figure S3**

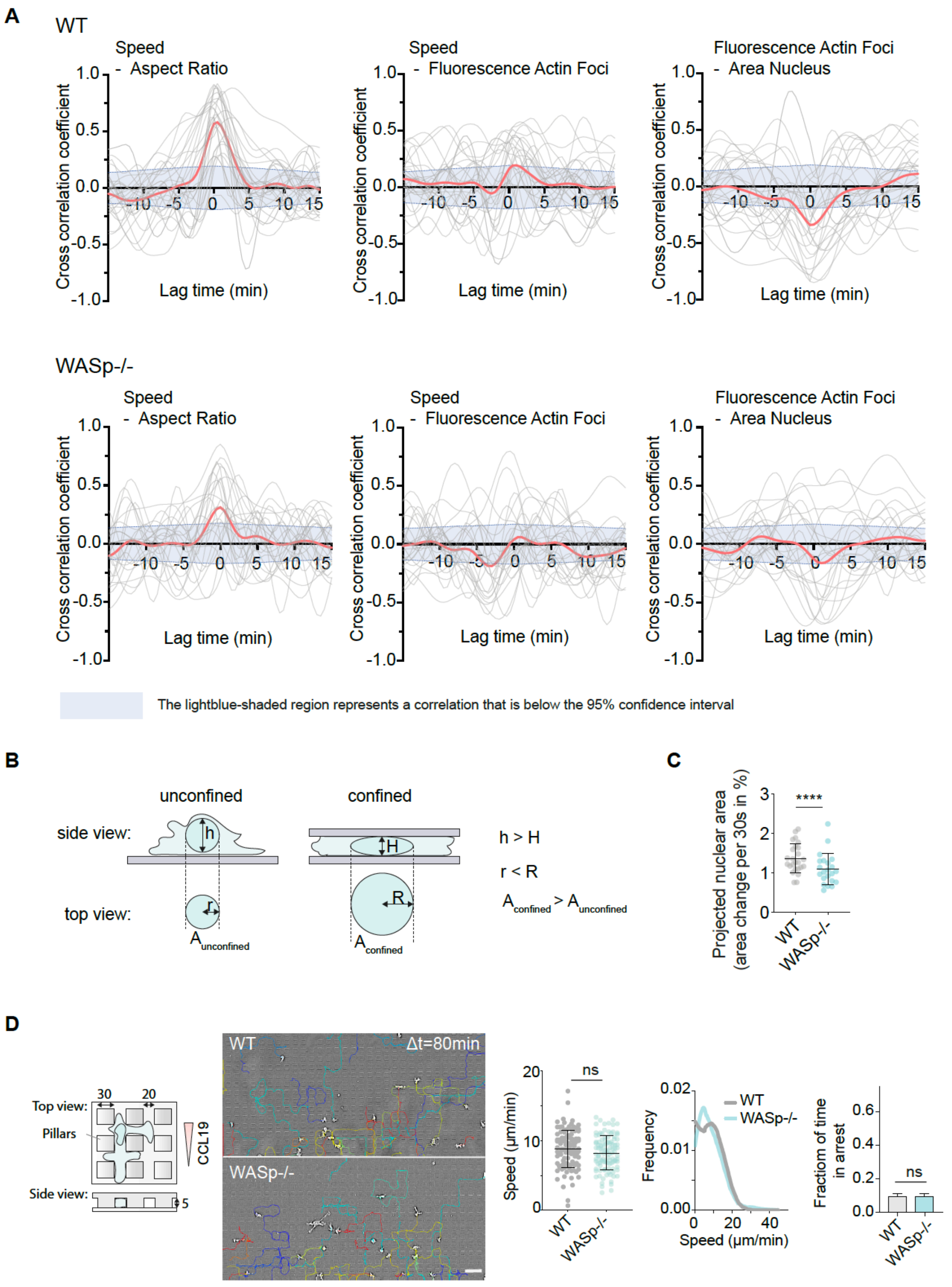
A
WT
Speed - Aspect Ratio
Speed - Fluorescence Actin Foci
Fluorescence Actin Foci - Area Nucleus
Cross correlation coefficient
Lag time (min)
WASp-/-
Speed - Aspect Ratio
Speed - Fluorescence Actin Foci
Fluorescence Actin Foci - Area Nucleus
The lightblue-shaded region represents a correlation that is below the 95% confidence interval
B
unconfined
confined
side view:
top view:
h > H
r < R
A confined > A unconfined
C
Projected nuclear area (area change per 30s in %)
****
WT
WASp-/-
D
Top view:
Pillars
Side view:
CCL19
WT
Δt=80min
WASp-/-
Speed (µm/min)
ns
Frequency
Speed (µm/min)
Fraction of time in arrest

**Figure S3. Related to Figure 3. Vertical pushing facilitates locomotion by deforming restrictive environments.** (**A**) Cross-correlation between the (1) Cell Speed and Aspect Ratio; (2) Cell speed and Mean Fluorescence of Actin Foci and (3) Mean Fluorescence of Actin Foci and Area of Nucleus. The thick red line represents the mean cross-correlation curve. Thin gray lines represent cross-correlation curves of individual cells (n=22-23). The light-blue shaded region represents a correlation that is below 95% confidence interval as obtained by bootstrapping. (**B**) Projected nuclear area as an indicator of cell height. The nucleus of unconfined DCs is approximately spherical (left), but naturally flattens with increasing confinement, making it a good indicator of cell height (right). h=height; r=radius; A=area. (**C**) Average change of nuclear area (in %) from one frame to the next frame (30s). Each data point shows the mean of an individual cell (movies were recorded for > 30 frames). Mann-Whitney test. **: p=0.0081. (**D**) DCs migrating in PDMS-based (non-deformable) microfluidic devices with a constant height. Left: Schematic showing dimensions of the pillar array. Representative brightfield micrograph from time-lapse movie showing DCs migrating in a non-deformable (PDMS) microfluidic device with pillars. Tracks of individual cells (color coded) observed for 80 min. Mean track speed: each data point represents one track; WT: n=110; WASp-/-: n=106 pooled from 3 experiments; Mann-Whitney test. Frequency distribution of instantaneous cell speeds (smoothed histogram). Fraction of time in arrest was extracted from tracks; Mean+SEM; Mann-Whitney test; ns=not significant.

### Figure S4

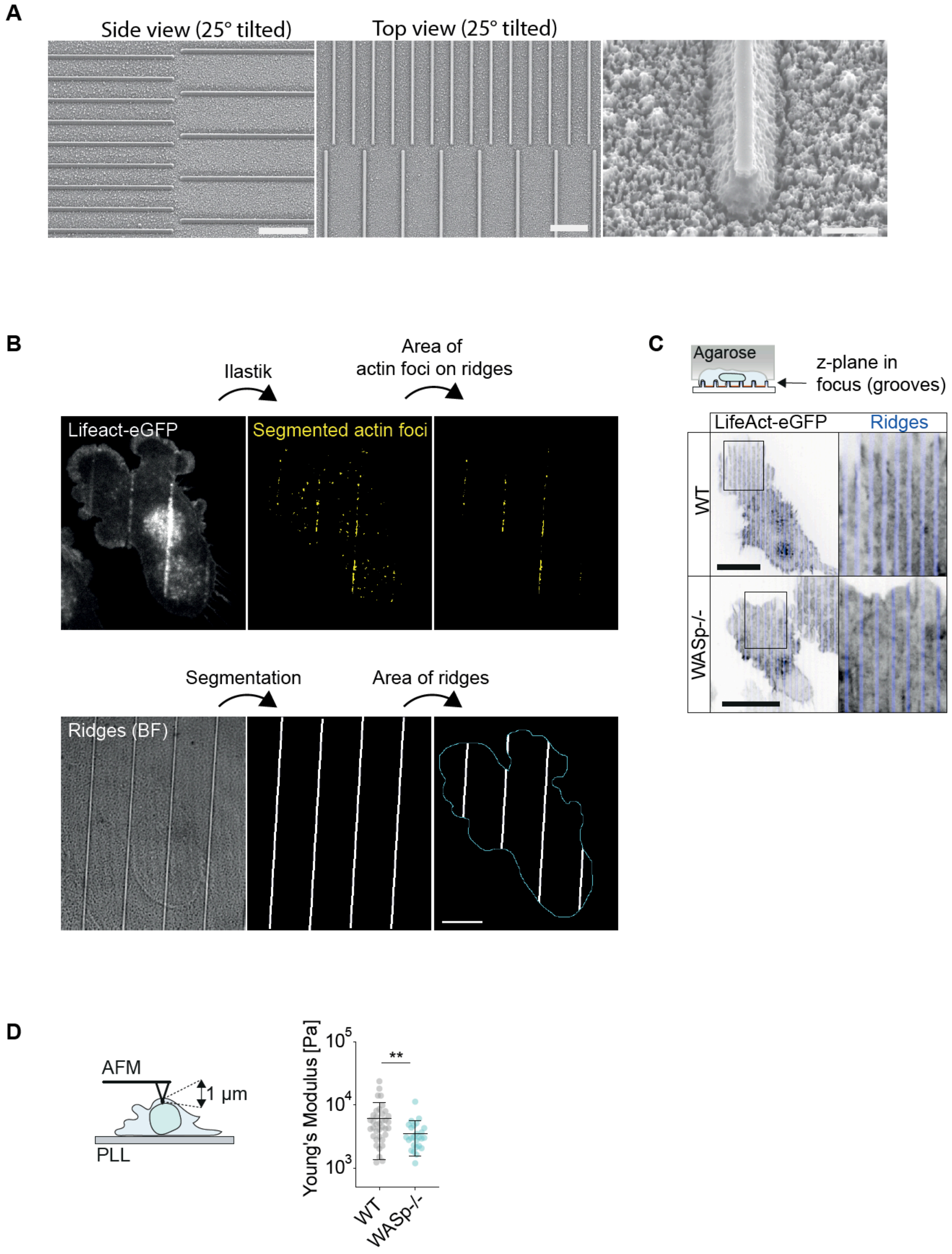
A
Side view (25° tilted)
Top view (25° tilted)
B
Ilastik
Area of
actin foci on ridges
Lifeact-eGFP
Segmented actin foci
Segmentation
Area of ridges
Ridges (BF)
C
Agarose
z-plane in
focus (grooves)
LifeAct-eGFP
Ridges
WT
WASp-/-
D
AFM
1 µm
PLL
Young's Modulus [Pa]
$10^5$
$10^4$
$10^3$
**
WT
WASp-/-

**Figure S4. Related to Figure 4. WASp-driven actin polymerization is triggered by mechanical loading.** (**A**) Scanning electron microscopy of coverslips with nano-ridges generated by electron beam lithography. Scale bars: left/middle=10 µm and right=1 µm. (**B**) Quantification of actin foci covering nano-ridges. Upper panel: Actin foci are segmented from LifeAct-eGFP micrographs (spinning disc microscopy) using Ilastik. Lower panel: Ridges are manually segmented from brightfield images using FIJI. Scale bar=10 µm. (**C**) Representative WT and WASp-/- DC migrating on nano-ridges. Focus on grooves. Notably, no actin foci are formed within grooves. Scale bar=20 µm. (**D**) Schematic of an AFM cantilever indenting a DC adherent to a PLL-coated coverslip (left). Using a Hertz contact mechanics model, the elastic modulus was estimated by fitting the force indentation curves up to 1 µm. Each data point represents one cell. WT: n=37 cells (also see Figure 1b); WASp-/-: n=26 cells; Mean±SD; Mann-Whitney test; **: p=0.0042.

### Figure S5

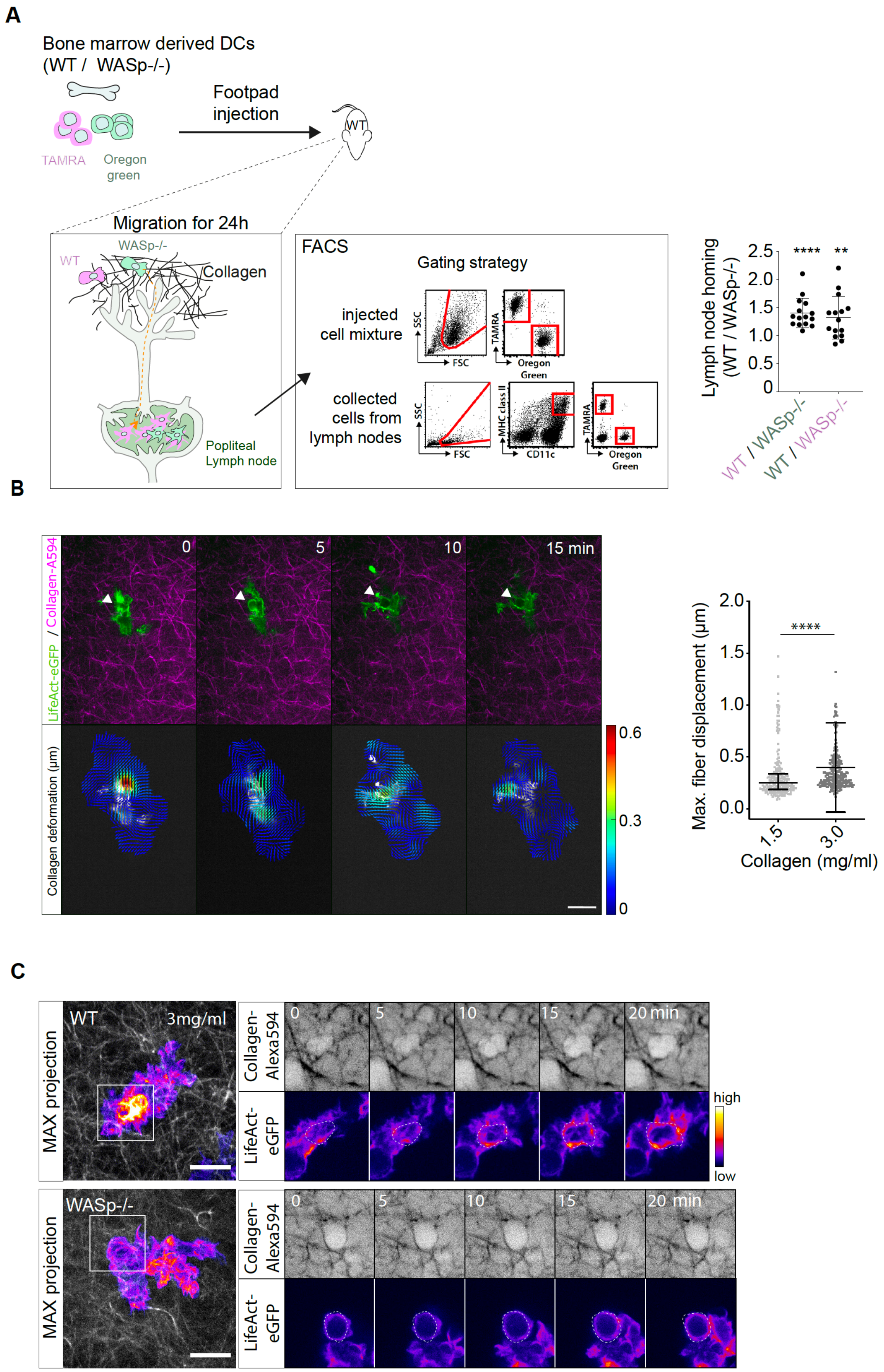
A
Bone marrow derived DCs
(WT / WASp-/-)
TAMRA
Oregon green
Footpad injection
WT
Migration for 24h
WT
WASp-/-
Collagen
Popliteal Lymph node
FACS
Gating strategy
injected cell mixture
collected cells from lymph nodes
SSC
FSC
TAMRA
Oregon Green
MHC class II
CD11c
Lymph node homing (WT / WASp-/-)
****
**
WT / WASp-/-
WT / WASp-/-
B
0
5
10
15 min
LifeAct-eGFP / Collagen-A594
Collagen deformation (μm)
0.6
0.3
0
Max. fiber displacement (μm)
Collagen (mg/ml)
1.5
3.0
C
WT
3mg/ml
WASp-/-
MAX projection
Collagen-Alexa594
LifeAct-eGFP
20 min
high
low

**Figure S5. Related to Figure 5. Deformation of collagen fibers is required for migration in fibrous environments.** (**A**) Left: Schematic drawing showing the workflow of lymph node homing experiments. Middle: Representative dot plots showing FACS gating strategy. Right: The mean ratio of WT DCs to WASp-/- DCs recruited to draining LNs was significantly larger than 1. Each data point represents the ratio of WT / WASp-/- of one experiment. To control for potential unspecific side effects of fluorescent labelling, fluorescent dyes were switched between WT and WASp-/-. WT (TAMRA) and WASp-/- (Oregon green): n=15; Ratio paired t-test; ****: p<0.0001; WT (Oregon green) and WASp-/- (TAMRA): n=15; Ratio paired t-test; **: p=0.0037. (**B**) Left: Time series of DC (LifeAct-eGFP) migrating in an Alexa-594-labeled collagen gel (3.0 mg/ml) (arrow head: collagen pore) (spinning disc confocal microscopy). Particle image velocimetry (PIV) of the same time series shows deformation of collagen fibers where a pore was dilated (arrow head). Displacement vectors are color coded for displacement in µm. Scale bar=15 µm. Right: Quantification: cells were imaged with 1 min frame-rate for 15-20 min. Each data point represents the maximum magnitude of frame-to-frame deformation; 1.5 mg/ml: n=245 data points pooled from 12 cells (3 independent experiments); 3.0 mg/ml: n=272 data points pooled from 14 cells (7 independent experiments); Median with interquartile range; Mann-Whitney test; ****=p<0.0001. (**C**) Representative time series of WT and WASp-/- DCs migrating in fibrous 3D collagen gels (3.0 mg/ml) (spinning disc confocal microscopy). Left panel shows maximum intensity projection of z scan. Scale bar=10 µm.

**Figure S6**

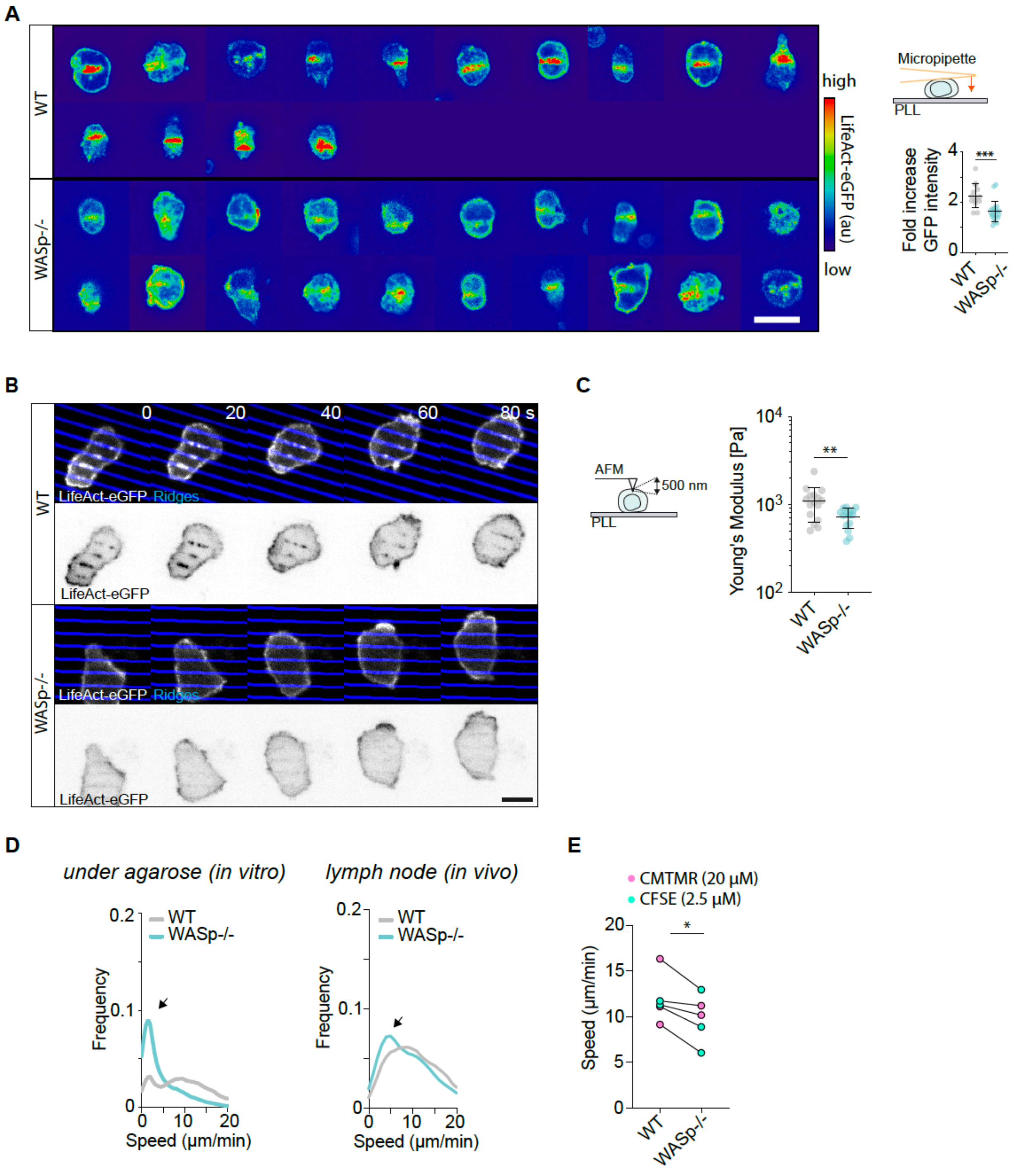
A
WT
WASp-/-
high
LifeAct-eGFP (au)
low
Micropipette
PLL
Fold increase GFP intensity
***
WT
WASp-/-
B
0
20
40
60
80 s
LifeAct-eGFP Ridges
LifeAct-eGFP
C
AFM
500 nm
PLL
Young's Modulus [Pa]
**
D
under agarose (in vitro)
lymph node (in vivo)
Frequency
Speed (µm/min)
E
CMTMR (20 µM)
CFSE (2.5 µM)
Speed (µm/min)
*

**Figure S6. Related to Figure 6. Orthogonal pushing drives T cell migration in crowded lymph nodes.** (**A**) T cells attached to poly-l-lysine (PLL)-coated coverslips were pinched from top with a microneedle (schematic). Representative micrographs (spinning disc confocal microscopy) of micropipette indentation experiments. LifeAct-eGFP intensity is color-coded. Scale bar=10 µm. Dot plot showing fold increase of LifeAct-eGFP intensity (area of indentation over non-indented area). Each data point is one indentation experiment. WT: n=14; WASp-/-: n=19; Mean±SD; Mann-Whitney test; ***: p=0.0002. (**B**) Representative time series (spinning disc confocal microscopy) of T cells migrating on nano-ridges. WT cells show sharply delineated actin foci matching the ridge structure. In contrast, formation of actin foci is abrogated in WASp-/- cells. Scale bar= 5 µm. (**C**) Schematic of an AFM experiment. Using a Hertz contact mechanics model, the elastic modulus was estimated by fitting the force indentation curves up to 500 nm. Each data point represents one cell. WT: n= 16; WASp-/-: n= 15; Mean±SD; Mann-Whitney test; **: p=0.0063. (**D**) Frequency distribution of instantaneous cell speeds (smoothed histogram). Left: under agarose (related to Figure 6A). Right: in lymph node (related to Figures 6F-H). (**E**) Pairwise analysis of differentially labeled WT and WASp-/- T cells migrating within the same LN. To control for potential unspecific side effects of fluorescent labelling, fluorescent dyes were switched between WT and WASp-/-. WT: n=5; WASp-/-: n=5; paired t-test; *: p= 0.0196.

### Figure S7

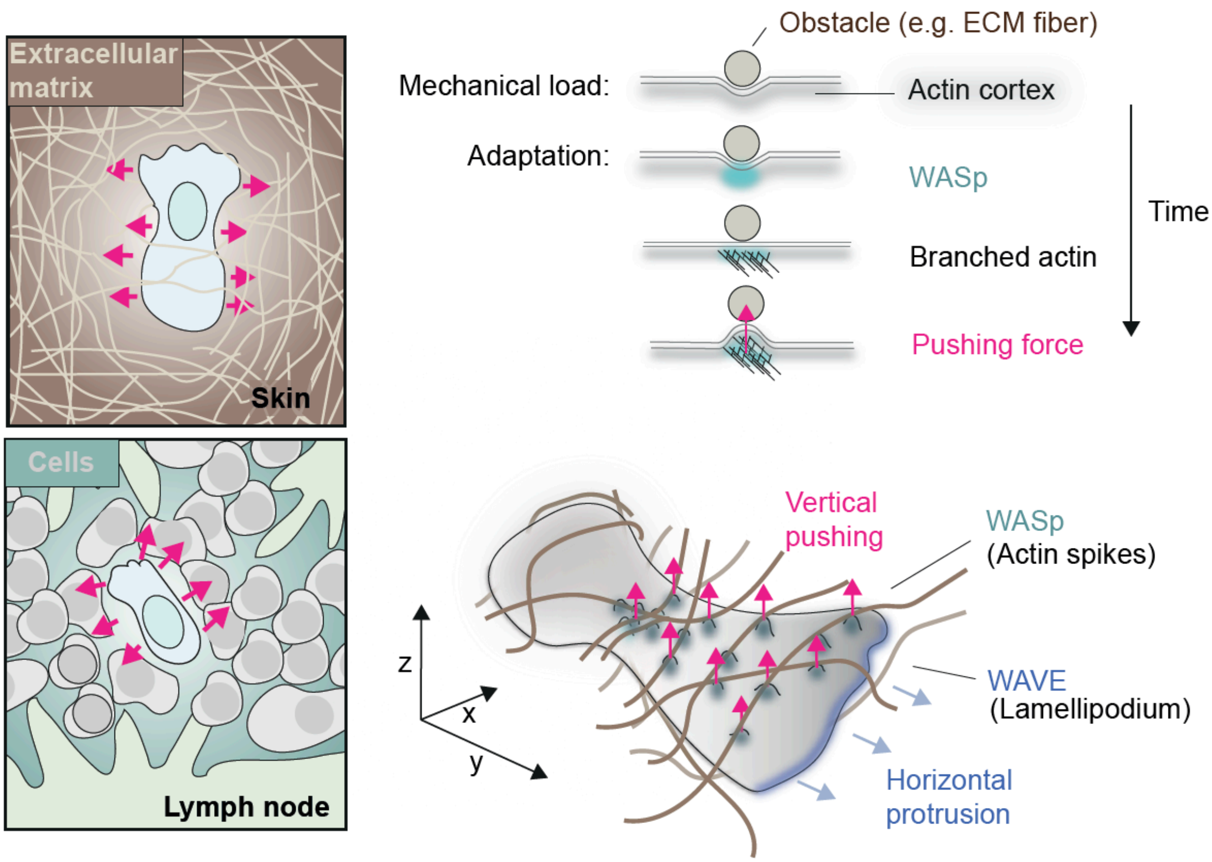


**Figure S7. Graphical summary: Migrating immune cells push their way through crowded tissues.** Right: Amoeboid migrating cells form actin spikes in response to the mechanical load of obstructive objects, such as collagen fibers. Local compression triggers formation of WASp foci that drive polymerization of branched actin networks. WASp-dependent actin networks are orthogonally orientated to WAVE-driven lamellipodia and push against restrictive obstacles to generate space for forward locomotion. Consequently, WASp is required for immune cell migration in crowded tissues such as skin and lymph nodes (left panels). Inherited loss-of-function of WASp is associated with immunodeficiency (Wiskott-Aldrich-Syndrom) and impaired immune cell migration.

### Supplementary Movie Legends

**Movie S1.**

Leukocytes migrating under mechanical load (related to Figures 1, S1). Epifluoresence movies (LifeAct-eGFP and Hoechst) of mature dendritic cells (DCs) migrating under agarose of low (2.5 kPa), intermediate (10 kPa) and high (17.5 kPa) stiffness. Lower panels: overview; upper panels: higher magnification.

**Movie S2.**

Dendritic cells form actin spikes in response to restrictive microenvironments (related to Figures 1, S1). (I) Spinning disc confocal microscopy of actin dynamics of migrating dendritic cells without confinement (2D), with partial confinement (3 um, 3D), with complete confinement (10 kPa agarose, 3D). (II) Dendritic cell migrating on inert substrate (PMOXA), confined under agarose. Retrograde movement of actin foci (spinning disc confocal microscopy). Actin foci were segmented in Ilastik and tracked with Trackmate. Tracks are color-coded (displacement).

**Movie S3.**

WASp polymerizes actin spikes and facilitates migration under confinement (related to Figures 2, S2). (I) WT, Hem1-/- and WASp -/- dendritic cells migrating under agarose (10 kPa); epifluorescence microscopy. Lower panels: overview; upper panels: hiher magnification. (II) TIRF microscopy of agarose confined DCs expressing Abi1-GFP (to visualize the WAVE complex) and WASp-eGFP (to visualize WASp). While Abi1-eGFP was strictly localized to the tip of lamellipodia, WASp-eGFP formed punctae scattered across the cell surface. (III) WASp drives actin spike formation scattered across the cell body while WAVE drives lamellipodia formation. Cells are confined under the load of agarose (10 kPa), on inert substrate (PMOXA) (Spinning disc microscopy). (IV) Epifluorescence movie of WT and WASp-/- DCs migrating under agarose. Actin Foci were segmented in Ilastik. Cell contours were stabilized and motion of actin foci is shown in the cell-frame of reference.

**Movie S4.**

Morphodynamic cycle of cellular locomotion under mechanical load (related to Figures 3, S3). (I) WT and WASp-/- dendritic cells migrating under the load of agarose (10 kPa); (Epifluorescence Microscopy). Right panels show color-coded LifeAct-eGFP signal. (II) Nuclear shape fluctuations of dendritic cells migrating under agarose. Dendritic cells show steady fluctuations of nuclear projected area when migrating under agarose (also see Figures S4B,C) (Epifluorescence microscopy).

**Movie S5.**

Dendritic cells migrating in non-deformable microfabricated PDMS devices (related to Figures 3, S3). (I) Migration of WASp-/- mutants in pillar forests is indistinguishable from WT cells (Brightfield microscopy). (II) WASp (WASp-eGFP) and F-actin (LifeAct-eGFP) are recruited to narrow constrictions (1-3 µm) (TIRF and brightfield microscopy).

**Movie S6.**

WASp-driven actin polymerization is triggered by mechanical loading (related to Figures 4, S4). (I) LifeAct-eGFP expressing WT and WASp-/- DCs were indented by the blunted end of a micropipette (Spinning disc microscopy). Last frame shows overlay with the outline of the micropipette. Lower panel shows color-coded LifeAct-eGFP signal. (II) LifeAct-eGFP expressing WT and WASp-/- DCs migrating on inert surfaces (PMOXA) with nano-ridges (Spinning disc microscopy). (III) WASp-eGFP expressing WT DCs migrating on inert surfaces (PMOXA) with nano-ridges (TIRF microscopy).

**Movie S7.**

Local actin polymerization pushes against restrictive collagen fibers (related to Figure 5). (I) Migrating dendritic cells (LifeAct-eGFP) deform collagen fibers (Alexa-594-labeled) (Spinning disc confocal microscopy). (II) WT DC (LifeAct-eGFP) migrating in a fibrous collagen gel (Alexa 594-labeled) (Spinning disc confocal microscopy). Local actin polymerization at sites of a narrow pore deforms restrictive collagen fibers thereby providing space for the cell body to move forward as indicated by shape changes of the nucleus (Hoechst).

**Movie S8.**

WASp-dependent actin foci drive t cell migration under mechanical load (related to Figures 6, S6). (I) Naïve T cells (CMTMR-labeled) migrating under agarose (10 kPa) (Epifluorescence microscopy). (II) Naïve T cells (LifeAct-eGFP) migrating under agarose (10 kPa) on inert (PLL-PEG) substrates. T cells show retrograde movement of actin foci

(Spinning disc confocal microscopy). Actin foci were segmented in Ilastik (show in yellow). (III) LifeAct-eGFP expressing WT and WASp-/- T cells were indented by the blunted end of a micropipette (Spinning disc microscopy). Lower panel shows overlay with the outline of the micropipette (LifeAct-eGFP is color coded). (IV) LifeAct-eGFP expressing WT and WASp-/- T-cells migrating on inert surfaces (PLL-PEG) with nano-ridges (Spinning disc microscopy).

**Movie S9.**

WASp drives T cell migration in crowded lymph nodes (related to Figure 6, S6). Adoptive transfer of fluorescently labeled T cells (WT = CMTMR (red) and WASp-/- = CFSE (green)) into wild-type recipient mice. After 24h, T cell migration in LN parenchyma was analyzed using intravital 2-photon microscopy.
